# Conformational biosensors delineate endosomal G protein regulation by GPCRs

**DOI:** 10.1101/2025.05.12.653522

**Authors:** Brian Wysolmerski, Emily E. Blythe, Mark von Zastrow

**Affiliations:** Department of Psychiatry and Behavioral Sciences, University of California, San Francisco, San Francisco CA, USA; Tetrad Graduate Program, University of California, San Francisco, San Francisco CA, USA; Department of Cellular and Molecular Pharmacology, University of California, San Francisco, San Francisco CA, USA

## Abstract

Many GPCRs trigger a second phase of G protein-coupled signaling from endosomes after initiating signaling from the plasma membrane. This inherently requires receptors to increase the concentration of active-state G proteins on the endosome membrane, but how this is achieved remains incompletely understood. We addressed this question by dissecting the regulation of G protein abundance and activity on endosomes following activation of several G_s_-coupled GPCRs–the β2-adrenergic receptor, the VIP-1 receptor, and the adenosine 2b receptor–that are natively co-expressed and differ in their ability to internalize after activation. We first verify GPCR-triggered redistribution of Gα_s_ from the plasma membrane to a mixed population of intracellular membranes, including endosomes, that is both reversible after receptor inactivation and triggered irrespective of the ability of the GPCR to internalize. We next show that GPCRs trigger this redistribution process at native expression levels and describe a method, using conformational biosensors, to detect endosomal activation of endogenous Gα_s_. Applying this method, we show that GPCR-mediated production of active-state Gα_s_ on endosomes depends on receptor endocytosis, whereas increasing the net amount of Gα_s_ on endosomes does not. Our results support a model for G_s_ regulation on endosomes mediated by two spatially separated receptor coupling events–one at the plasma membrane controlling endosomal G_s_ abundance and another at endosomes controlling G_s_ activity. Additionally, our results reveal location-bias in the selectivity of G protein activation on endosomes that is differentially programmed by GPCRs in a receptor-specific manner.

## Introduction

G protein-coupled receptors (GPCRs) constitute the largest family of signaling receptors that regulate nearly every physiological process. After activation by binding an agonist, GPCRs initiate signaling by coupling to cognate heterotrimeric G proteins, consisting of a Gα subunit and Gβγ subcomplex, which function as key transducers of downstream signaling. This allosteric coupling reaction promotes guanine nucleotide exchange on the G protein α-subunit, resulting in GTP binding to the α-subunit that converts it from an inactive to active state. G protein classes are defined according to the identity of their α-subunit (e.g. G_s/olf_, G_i/o_, G_q/11_, and G_12/13_), with individual GPCRs differing in selectivity for coupling among G protein classes, and each class producing distinct downstream regulatory effects^1,2^. The central importance of GPCR signaling in physiology, and its dysfunction or dysregulation in pathological states, has motivated intense interest in GPCRs as therapeutic targets, with over 30% of presently FDA-approved drugs targeting GPCRs^3^.

The importance of GPCR signaling from the plasma membrane has been recognized for many years^1,4^, and there is now considerable interest in the ability of GPCRs to produce distinct and additional effects from intracellular membranes^4,5^. Endomembrane signaling is perhaps most strongly supported from the study of G_s_-coupled GPCRs^4–16^, but there is also significant evidence for endomembrane signaling through other G protein classes^5,17,18^. Such signaling fundamentally depends on the activated GPCR increasing the concentration of active-state G proteins on the appropriate membrane. When compared to the present knowledge about the subcellular localization of activated GPCRs, however, relatively little is known about how active-state G protein concentration on endomembranes is controlled.

G_s_ is enriched on the plasma membrane and present in lower amounts on intracellular membranes^19^. G_s_ activation by coupling to a GPCR on the plasma membrane promotes dissociation of Gα_s_ and its net intracellular redistribution, increasing the concentration of Gα_s_ on multiple endomembrane compartments, including endosomes^19–23^. Producing active-state

Gα_s_ on endosomes is then thought to require a second coupling reaction locally on endosomes^4,5,9^. However, the presence of active-state Gα_s_ on endosomes has only been hypothesized, and the cellular basis of the regulation of Gα_s_ abundance and activity on endosomes remains unclear.

We addressed this knowledge gap by dissecting the regulation of endosomal Gα_s_ abundance and activity by GPCRs that are natively co-expressed and significantly differ in their ability to internalize after activation. We first verify that GPCRs trigger a net intracellular redistribution of Gα_s_ from the plasma membrane^19,20,22–24^, and we then extend the present understanding by showing that this process is triggered by multiple G_s_-coupled GPCRs and occurs under conditions of native or near-native levels of GPCR and Gα_s_ expression. Next, using conformational biosensors, we demonstrate sequential phases of both G_s_ activation and of active-state Gα_s_ accumulation, first on the plasma membrane and then endosomes, at endogenous levels of G protein expression. We then show that G_s_ activation on endosomes is specifically dependent on receptor endocytosis. Finally, we provide evidence for a type of ‘location bias’ in the biochemical selectivity of endosomal G protein activation that is differentially programmed by GPCRs in a remarkably receptor-specific manner.

## Results

### Gα_s_ colocalizes with internalized receptors and Gβγ on early endosomes after GPCR activation

A prerequisite for GPCRs to increase the concentration of active-state Gα_s_ on endosomes is for Gα_s_ to be present on the relevant endosome membrane. It is generally thought that Gα_s_ dissociates from the plasma membrane after its activation and subsequently samples a variety of intracellular membrane compartments, including endosomes that contain activated GPCRs after regulated endocytosis^19,22,23,25,26^. A second round of GPCR-triggered G protein activation is then thought to occur on endosomes^4,5,7,9,10^. We sought to verify and further examine this process in living cells. We began by imaging Gα_s_ in living cells by confocal microscopy, using the β2-adrenergic receptor (β2AR) as a model G_s_-coupled GPCR that is well known to trigger intracellular redistribution of Gα_s_^19,20,22–24^. We labeled Gα_s_ by inserting EGFP into the linker region between its α-helical appendage and conserved Ras-like domain, a strategy shown previously to preserve the signaling function of Gα_s_^24^, and we then expressed this labeled construct in HEK293 cells stably expressing Flag-tagged β2ΑR. Consistent with previous studies from others and us^20,22–24^, EGFP-Gα_s_ visibly redistributed intracellularly from the plasma membrane within several minutes after application of the β2AR agonist, isoproterenol (Iso, Fig. 1a). EGFP-Gα_s_ was diffusely distributed in the cytoplasm and concentrated on various endomembranes, including endosomes, as verified by colocalization with mApple-EEA1 (Supplementary Fig. 1a). We previously observed redistribution from the plasma membrane to the cytoplasm, but we were unable to resolve specific internal membrane localization^20^. As these early studies were conducted using epitope-tagged Gα_s_ in fixed cells, we examined fixed cells and found that intracellular membrane localization of EGFP-Gα_s_ was less well-preserved (Supplementary Fig. 1b).

**Figure 1:**
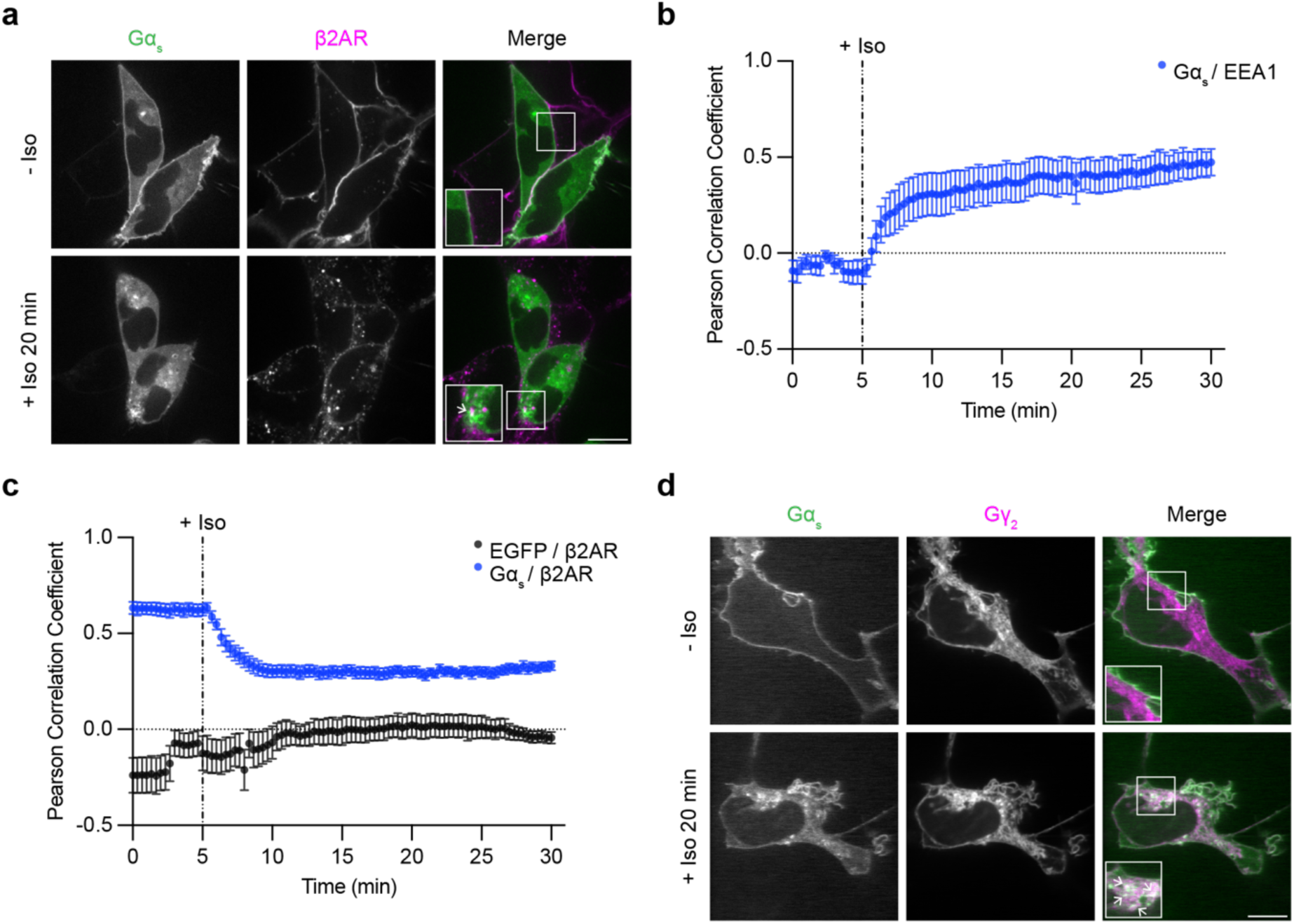
Gα_s_ colocalizes with internalized β2ΑRs and Gβγ on endosomes after receptor activation. **a)** Representative confocal images of live HEK293 cells stably expressing Flag-β2AR and transiently expressing EGFP-Gα_s_ before and after 20 minutes of Iso (1 µM) treatment. **b)** Pearson correlation coefficient between Gα_s_ and mApple-ΕΕΑ1 channels over time. Images are shown in Supplementary Fig. 1a. **c)** Pearson correlation coefficient between Gα_s_ and either the Flag-β2AR or EGFP channels over time. Significance determined by repeated measures 2-way ANOVA with Sidak’s multiple comparisons test (see Supplementary Table 4 for p values). **d)** Representative confocal images of live HEK293 cells stably expressing Flag-β2AR and transiently expressing EGFP-Gα_s_ and mApple-Gγ_2_ before and after 20 minutes of Iso (1 µM) treatment. Images are representative of at least 3 independent experiments. Prior to imaging, cells were treated for 10 minutes with an anti-Flag antibody coupled to Alexa Fluor 647 to label surface Flag-β2AR. In panel a, cells were co-transfected with myc-Gβ_1_ and untagged Gγ_2_. In panel d, cells were cotransfected with untagged Gβ_1_. Scale bars = 10 μm. Insets in panels a and d are 1.5x zoom of indicated regions and arrows indicate examples of colocalization. For Pearson correlation analysis, data are represented as mean ± S.E.M. of individual dishes (n = 6-16) from at least 3 independent experiments. Iso (1 µM) was added after 5 minutes of imaging, depicted by the dashed line.

Gα_s_ activation on endosomes would require receptors to be present in the same endosome membrane. Previous studies have reported little or no colocalization between Gα_s_ and internalized β2ARs^22,23^, so we tested this in our system by imaging EGFP-Gα_s_ and internalized β2ARs. Confocal microscopy resolved EGFP-Gα_s_ localization on Flag-β2AR-containing endosomes in isoproterenol-treated cells, and we also observed EGFP-Gα_s_ localization on additional compartments not containing internalized receptors (Fig. 1a, arrows indicate examples of EGFP-Gα_s_ / Flag-β2AR colocalization). We quantified colocalization using Pearson correlation analysis, assessing pixel-based correlations between EGFP-Gα_s_ and either the mApple-EEA1 or Flag-β2AR fluorescence signal, respectively, after agonist application. The Pearson correlation coefficient between EGFP-Gα_s_ and mApple-EEA1 increased after receptor activation (Fig. 1b), indicating Gα_s_ redistribution to early endosomes. In contrast, the Pearson correlation coefficient between EGFP-Gα_s_ and Flag-β2AR decreased after agonist application (Fig. 1c), likely representing loss of both Flag-β2AR and EGFP-Gα_s_ from the plasma membrane. However, the Pearson correlation coefficient between EGFP-Gα_s_ and Flag-β2ΑR plateaued at a level that remained significantly higher than the Pearson correlation between a cytosolic EGFP control and Flag-β2ΑR (Fig. 1c), consistent with our observations of partial colocalization with receptors on endosomes. Together, these results indicate that activated β2ARs trigger EGFP-Gα_s_ to rapidly redistribute from the plasma membrane to endomembranes, including endosomes that also contain internalized β2ARs.

As Gα_s_ association with Gβγ is a prerequisite for the GPCR coupling reaction that produces active Gα_s_, we asked if Gβγ is present on the same endosomes using a labeled γ-subunit (mApple-Gγ_2_) to detect Gβγ. Labeled Gβγ was broadly distributed on the plasma membrane and on multiple internal membranes, including membranes also associated with ΕGFP-Gα_s_ (Fig. 1d). Whereas endosomal localization of labeled Gα_s_ and β2AR were both markedly increased after β2AR activation, the localization of Gβγ appeared similar both before and after application of isoproterenol (Fig. 1d). These findings support a model in which Gα_s_ and β2AR colocalize on endosomes in an activation-induced manner, and that these endosomes are constitutively associated with Gβγ^27^. Accordingly, all of the protein components necessary for G protein coupling are present on endosomes after agonist-induced activation.

### Gα_s_ returns to the plasma membrane after receptor inactivation

The ability of cells to respond to a subsequent agonist exposure would presumably require replenishment of Gα_s_ at the plasma membrane, and previous studies have suggested that the intracellular redistribution of Gα_s_ triggered by β2AR activation is reversible after receptor inactivation^19,20^. We sought to verify this by activating Flag-β2ΑR with Iso in cells coexpressing EGFP-Gα_s_, and then applying the β2AR antagonist Alprenolol (Alp) in excess to inactivate the receptor. We observed a pronounced reaccumulation of EGFP-Gα_s_ on the plasma membrane after β2AR inactivation (Fig. 2a, Supplementary Fig. 2a), verifying reversibility of the Gα_s_ redistribution process as reported previously by others and us^19,20^. To quantify these effects, we used NanoBit protein complementation to measure the plasma membrane localization of Gα_s_ over time. We inserted LgBit into Gα_s_ at the same position as EGFP (LgBit-Gα_s_) and measured complementation with a plasma membrane-targeted SmBit construct (SmBit-mApple-CAAX), confirming appropriate plasma membrane localization of this construct by confocal microscopy (Supplementary Fig. 2b). In this assay, a net intracellular redistribution of Gα_s_ results in a decrease in the luminescence signal, which is produced by protein complementation (Fig. 2b). Iso-induced activation of Flag-β2AR produced a decrease in luminescence intensity with a similar time course as the redistribution observed by microscopy (Fig. 2c), and this signal recovered to near baseline after reversal of β2AR activation (Fig. 2c). These data verify that net intracellular redistribution of Gα_s_ triggered by β2AR activation is reversible after receptor inactivation, resulting in a net replenishment of the plasma membrane-associated Gα_s_ pool.

**Figure 2:**
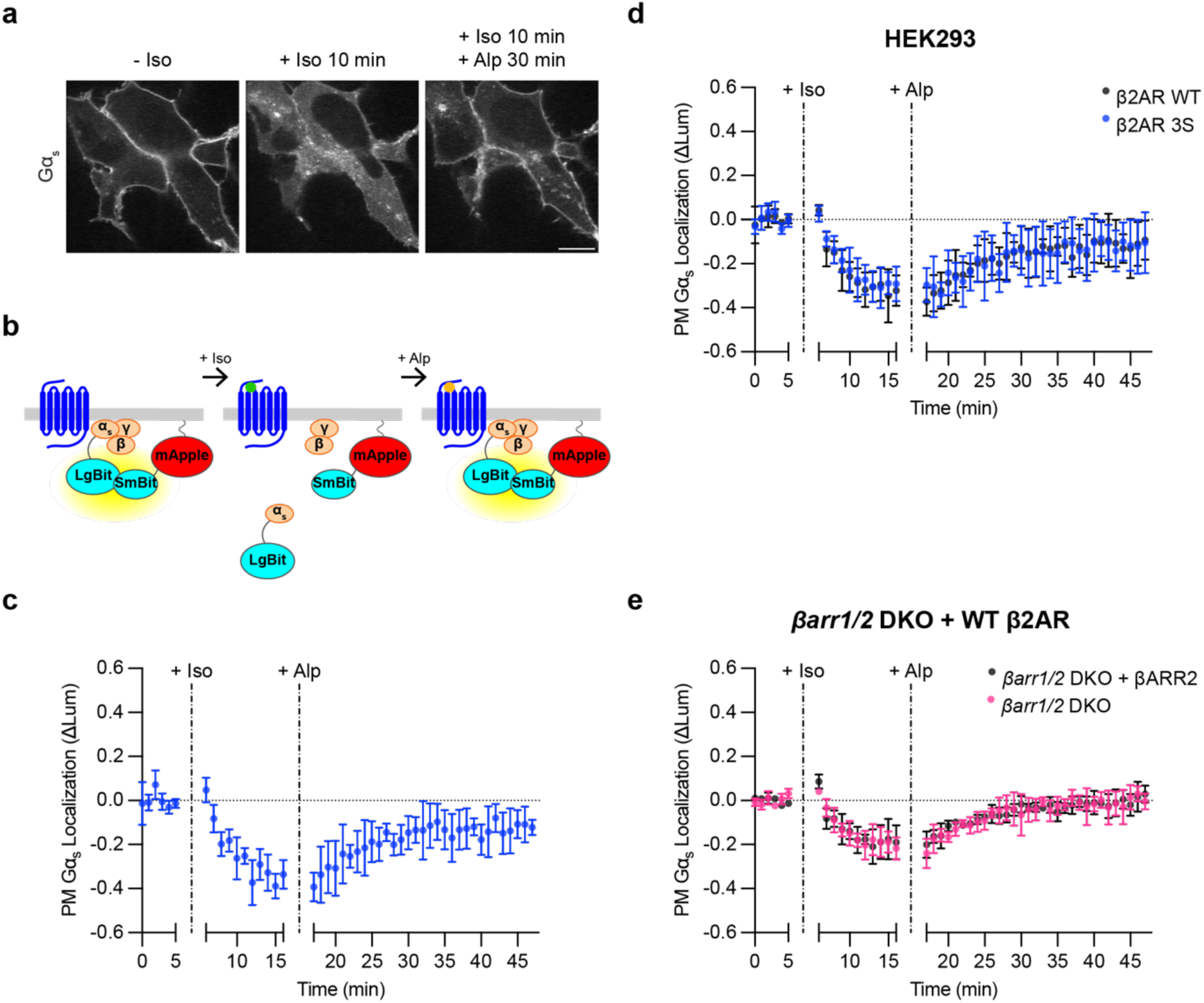
Redistribution of Gα_s_ is reversible after receptor inactivation and independent of receptor endocytosis. **a)** Representative stills from time-lapse confocal microscopy of live HEK293 cells stably expressing Flag-β2AR and transfected with EGFP-Gα_s_, myc-Gβ_1_, and untagged Gγ_2_ either before drug treatment, after 10 minutes of Iso (100 nM) treatment, or after 10 minutes of Iso followed by 30 minutes of Alprenolol (Alp, 10 μΜ) treatment. Images are representative of 4 independent experiments. Scale bar = 10 μm. **b)** Schematic of plasma membrane Gα_s_ NanoBit bystander assay. **c)** NanoBit bystander assay showing plasma membrane localization of Gα_s_ in HEK293 cells expressing WT Flag-β2AR after Iso (100 nM at 5 minutes) followed by Alp (10 µM at 16 minutes) treatment. **d)** NanoBit bystander assay showing plasma membrane localization of Gα_s_ after Iso (100 nM at 5 minutes) treatment followed by Alp (10 µM at 16 minutes) treatment in HEK293 cells expressing either Flag-β2ΑR WT or Flag-β2ΑR-3S. Significance (n.s.) determined by two-way ANOVA with Sidak’s multiple comparisons test (see Supplementary Table 5). **e)** NanoBit bystander assay showing plasma membrane localization of Gα_s_ after Iso (100 nM at 5 minutes) followed by Alp (10 µM at 16 minutes) treatment in *βarr1/2* DKO HEK293 cells expressing either βARR2-mApple or mApple. Significance (n.s.) determined by repeated measures two-way ANOVA with Sidak’s multiple comparisons test (see Supplementary Table 6). Data are shown as mean ± S.D. of at least 3 biological replicates.

### Intracellular redistribution of Gα_s_ does not depend on GPCR endocytosis

Previous studies differ in whether intracellular redistribution of Gα_s_ triggered by β2ARs requires receptor endocytosis^20,22,23^, so we revisited this question using improved tools for monitoring Gα_s_ redistribution. As a first approach, we blocked endocytosis of β2ARs using the dynamin inhibitor Dyngo4a^28^. Iso-induced internalization of Flag-β2AR was strongly suppressed by Dyngo4a treatment; however, intracellular redistribution of EGFP-Gα_s_ was not detectably affected, suggesting that endocytosis of the activating GPCR is not required for GPCR-triggered intracellular redistribution of Gα_s_ (Supplementary Fig. 3a).

As a second approach, we prevented receptor internalization by mutating specific serine residues in the cytoplasmic tail of β2AR to alanine^9,29,30^, and we assessed the ability of the internalization-defective mutant receptor to trigger intracellular redistribution of Gα_s_. HEK293 cells endogenously express β-adrenergic receptors at a low level^31^, but this level was not sufficient to produce a detectable Iso-induced redistribution of Gα_s_ in the NanoBit assay (Supplementary Fig. 3b). Thus, we compared the effects of β2AR-3S to WT β2AR when overexpressed at comparable levels (Supplementary Fig. 3c). β2AR-3S triggered intracellular redistribution of Gα_s_ that was indistinguishable from that triggered by WT β2AR (Fig. 2d), and we verified by imaging that β2AR-3S also promoted accumulation of EGFP-Gα_s_ on endomembranes (Supplementary Fig. 3d).

Finally, as a third approach, we examined Gα_s_ redistribution in previously described β-arrestin 1/2 double knockout (*βarr1/2* DKO) HEK293 cells^15^. After verifying a blockade of WT β2AR internalization (Supplementary Fig. 3e), we again observed that Iso treatment triggered robust and reversible intracellular redistribution of Gα_s_ in these cells (Fig. 2e, Supplementary Fig. 3e). Moreover, re-expression of βARR2-mApple in this knockout background fully rescued internalization of Flag-β2ΑR but did not affect the observed redistribution of Gα_s_ (Fig. 2e, Supplementary Fig. 3e). Together, these results strongly suggest that receptor internalization is not required for the intracellular redistribution of Gα_s_.

*Intracellular redistribution of Gα_s_ is triggered by multiple G_s_-coupled GPCRs at native levels* We next asked if the ability to trigger intracellular redistribution of Gα_s_ is shared by other G_s_-coupled GPCRs. We focused on the adenosine-2B receptor (A_2B_R or ADORA2B) and vasoactive intestinal peptide-1 receptor (VIPR1 or VPAC1) as representatives of GPCR families A and B, respectively, each of which is natively expressed in HEK293 cells^31^. We found both receptors triggered a pronounced intracellular redistribution of EGFP-Gα_s_ when overexpressed and activated by their cognate agonists (VIP or NECA), similar to that triggered by overexpressed β2AR after activation by Iso (Supplementary Fig. 4).

Because achieving GPCR-triggered redistribution of EGFP-Gα_s_ in these assays required the receptor to be overexpressed, we wondered if Gα_s_ redistribution can also be triggered by native GPCRs or, alternatively, if this redistribution process is an artifact of excessive receptor expression. We reasoned that overexpressing EGFP-Gα_s_ in cells also expressing endogenous, unlabeled Gα_s_ might limit the sensitivity of our assay for detecting redistribution triggered by endogenous receptors, particularly as the level of native receptor expression is far lower than that of Gα_s_^32^. To address this, we expressed EGFP-Gα_s_ at a much lower level in cells lacking native Gα_s_, which we reasoned would likely compete in our assay. We first knocked out endogenous Gα_s_ by CRISPR-Cas9 editing of the *GNAS* gene (Gα_s_ KO cells).

Full knockout of Gα_s_ was verified genetically and by immunoblot, as well as functionally by loss of the Iso-induced cytoplasmic cAMP elevation (Supplementary Fig. 5). We then isolated cell populations stably expressing EGFP-Gα_s_ at a low level, within 2-fold of the native level in wild-type cells as assessed by immunoblot, and we verified rescue of the Iso-induced cAMP response in two independent isolates (Supplementary Fig. 6).

This near-native level of EGFP-Gα_s_ expression was detectable by confocal microscopy but quite dim (Supplementary Fig. 6c), so we turned to total internal reflection fluorescence (TIRF) microscopy as a more sensitive and quantitative method for assessing EGFP-Gα_s_ localization. We detected intracellular redistribution of EGFP-Gα_s_ by loss of fluorescence intensity from the evanescent field, in which the bottom plasma membrane surface is selectively illuminated relative to the cell interior. Using this approach, we observed a significant Iso-induced reduction of surface-localized EGFP-Gα_s_ upon stimulation of endogenous β2ARs with Iso, indicating a net intracellular redistribution of EGFP-Gα_s_ from the plasma membrane. The magnitude of this effect was much lower than that observed in the presence of overexpressed β2ARs (Fig. 3 and Supplementary Fig. 7, Supplementary Tables 8 and 9), consistent with a much lower level of endogenous receptor expression.

**Figure 3:**
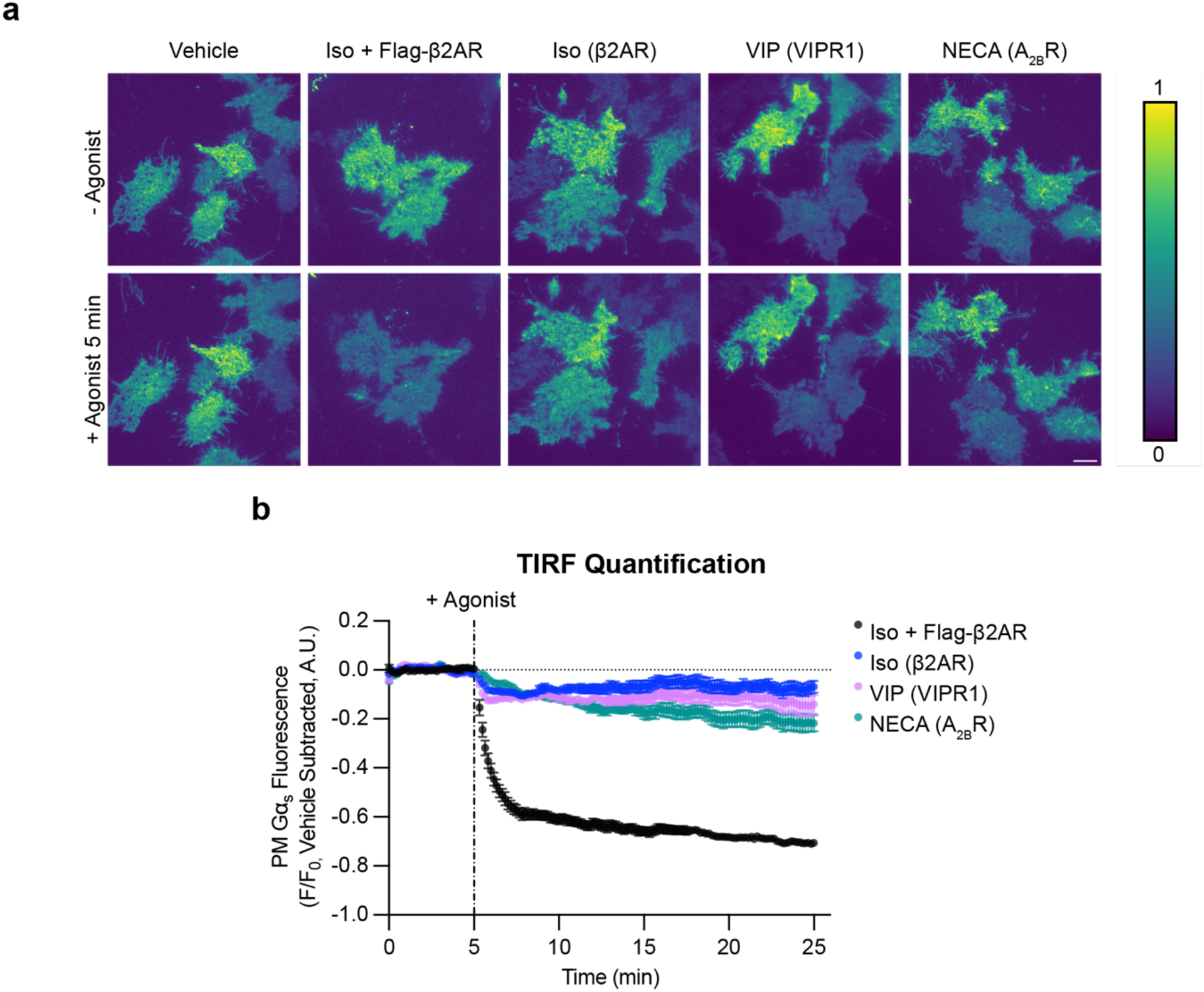
Activation of 3 endogenously expressed GPCRs drives Gα_s_ redistribution from the plasma membrane. **a)** Representative stills from time-lapse TIRF microscopy of Gα_s_ KO1 + EGFP-Gα_s_ rescue cells stably expressing EGFP-Gα_s_ before or after 5 minutes of agonist treatment to activate endogenously expressed GPCRs (β2AR, VIPR1, A_2B_R). Images are shown as heat maps normalized to t = 3 minutes before drug addition for each individual movie. Cells were treated with either vehicle, Iso (1 µM, ± overexpression of Flag-β2AR), VIP (1 µM), or NECA (20 µM). Scale bar = 10 µm. **b)** Quantification of Gα_s_ fluorescence from TIRF movies depicted in a). The average of vehicle control movies (n = 5) at each time point was subtracted before plotting data. n ≥ 4 movies from at least 2 experiments. Data are represented as mean ± S.E.M. of individual movies. Significance determined by repeated measures 2-way ANOVA with Dunnett’s multiple comparisons test (see Supplementary Table 8 for p values).

Both VIP and NECA also triggered a significant intracellular redistribution of EGFP-Gα_s_ through their endogenously expressed receptors (Fig. 3 and Supplementary Fig. 7, Supplementary Tables 8 and 9). Together, these results indicate that a variety of G_s_-coupled GPCRs share the ability to trigger the intracellular redistribution of Gα_s_, and that they can do so at native or near-native levels of receptor and G protein expression.

### VIPR1 produces active-state Gα_s_ on the plasma membrane and endosomes

Signaling from endosomes requires Gα_s_ associated with the endosome membrane to be in an active state. The above experiments provided useful information about the net subcellular redistribution of Gα_s_, but they provided no information about its activation state. To address this, we utilized KB1691, a peptide that binds selectively to active-state Gα_s_ and has been used previously as a biosensor for detecting active-state Gα_s_ in cells that overexpress both receptors and G_s_^33^. We wondered if this peptide can be adapted to resolve the location of active-state Gα_s_ at a native expression level. We tested this by focusing on VIPR1 because this GPCR strongly stimulates sequential G_s_ signaling phases from the plasma membrane and endosomes^15^. We began by assessing VIP-stimulated recruitment of KB1691 to the plasma membrane, fusing KB1691 to SmBit-mApple (SmBit-mApple-KB1691) and measuring protein complementation with a plasma membrane-targeted LgBit (LgBit-CAAX, Fig. 4a). A clear VIP-induced recruitment signal was detected in cells expressing only native G proteins (Fig. 4b), albeit dependent on VIPR1 being overexpressed. These results verify the utility of KB1691 as a biosensor and indicate that KB1691 is sufficiently sensitive to detect active-state G proteins at native levels in cells that overexpress the activating GPCR.

**Figure 4:**
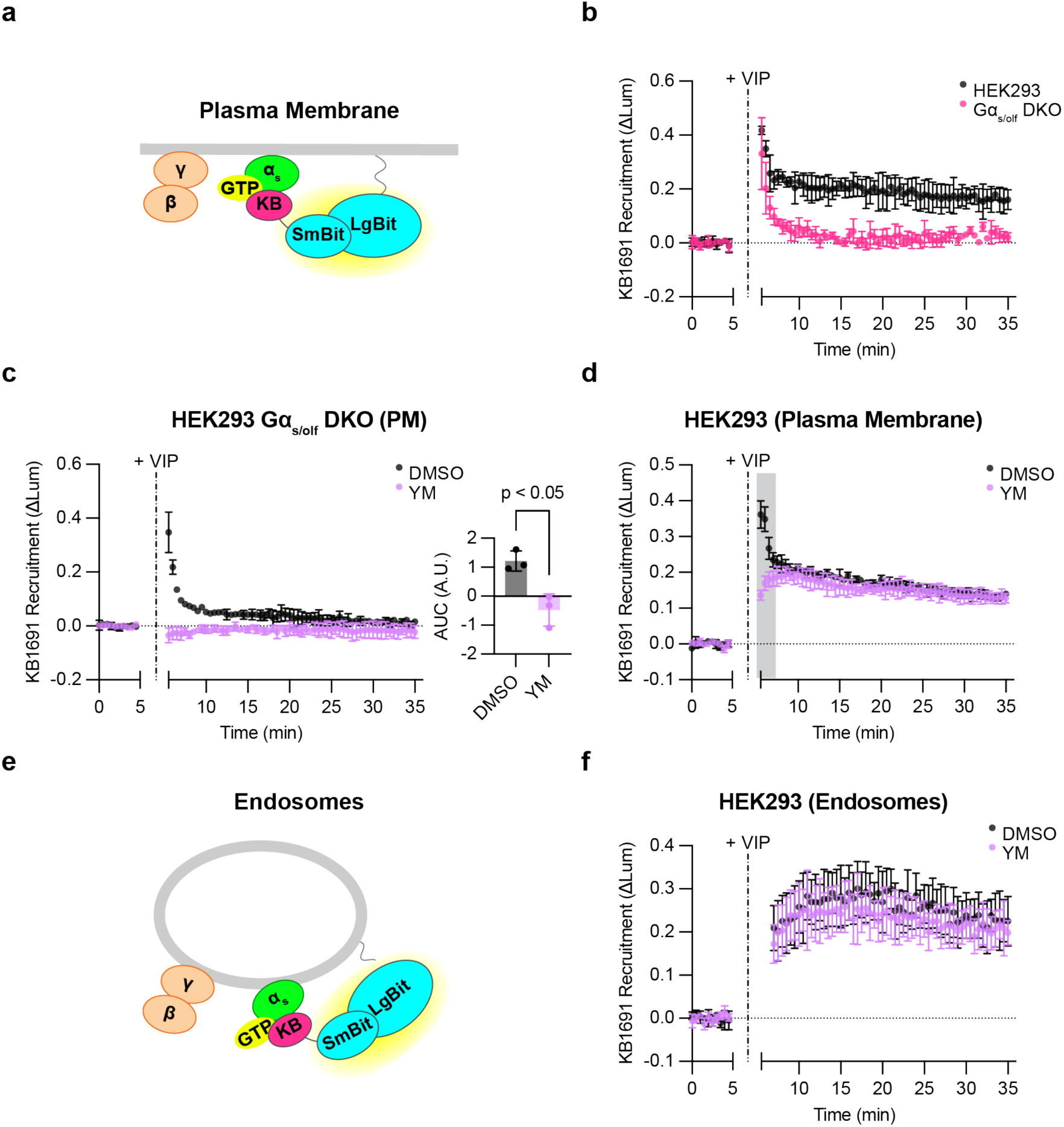
Detection of active, endogenous G proteins after VIPR1 activation. **a)** Schematic of KB1691 (active-state Gα_s_ biosensor) plasma membrane NanoBit bystander assay. **b)** NanoBit bystander assay showing recruitment of KB1691 to the plasma membrane in both HEK293 parental cells and Gα_s/olf_ DKO cells expressing Halo-VIPR1. VIP (1 µM) was added after 5 minutes. **c)** Left: NanoBit bystander assay showing recruitment of KB1691 to the plasma membrane in Gα_s/olf_ DKO cells expressing Halo-VIPR1 and pretreated with either DMSO (0.1 %) or YM-254890 (1 µM, 30 minutes). Right: AUC of time course. VIP (1 µM) was added after 5 minutes. Significance determined by two-tailed unpaired t test. **d)** NanoBit bystander assay showing recruitment of KB1691 to the plasma membrane in HEK293 cells expressing Halo-VIPR1 and pretreated with either DMSO (0.1 %) or YM-254890 (1 µM, 30 minutes). VIP (1 µM) was added after 5 minutes. Shaded areas represent time points at which the difference between DMSO- and YM-treated cells are statistically significant (p < 0.05, determined by repeated measures 2-way ANOVA with Sidak’s multiple comparisons test, see Supplementary Table 10). **e)** Schematic of KB1691 endosome NanoBit bystander assay. **f)** NanoBit bystander assay showing recruitment of KB1691 to endosomes in HEK293 cells expressing Halo-VIPR1 and pretreated with either DMSO (0.1 %) or YM-254890 (1 µM, 30 minutes). VIP (1 µM) was added after 5 minutes. Significance (n.s., see Supplementary Table 11) determined by repeated measures 2-way ANOVA with Sidak’s multiple comparisons test. Data are shown as mean ± S.D. of at least 3 independent experiments.

To test the specificity of sensor recruitment, we used knockout cells lacking Gα_s_, and additionally deleted Gα_οlf_, a close paralog of Gα_s_ (Gα_s/olf_ DKO cells) that we thought might also engage the sensor (Supplementary Fig. 5). The VIP-induced biosensor response was markedly reduced in Gα_s/olf_ DKO cells, verifying the ability of the sensor to detect native Gα_s_ activity in our assay (Fig. 4b). However, we were surprised to observe a residual signal remaining in Gα_s/olf_ DKO cells that exhibited faster onset and shorter duration (Fig. 4b).

Further, we verified in the same DKO cell background that the VIP-induced cAMP response was abolished, and that it was rescued by EGFP-Gα_s_ re-expression as expected (Supplementary Fig. 5 and 6). Together, these kinetic and genetic data strongly suggest that the sensor detects both active-state Gα_s_ and a distinct, non-G_s_ component.

As VIPR1 has been observed previously to couple to G_q_ as well as G_s_^34^, we wondered if this residual signal might represent active-state Gα_q_ and/or Gα_11_. Consistent with this possibility, the G_q/11_ inhibitor YM-254890 (YM)^35^ eliminated the residual VIP-induced recruitment signal in Gα_s/olf_ DKO cells (Fig. 4c). Furthermore, YM selectively blocked the rapid component of biosensor recruitment measured in wild type cells without affecting the slower component (Fig. 4d). Accordingly, in subsequent experiments, we defined the biosensor signal measured in the presence of YM as a specific readout of active-state G_s_. We interpret the YM-sensitive fraction of the signal simply as a distinct, non-G_s_ component, which may represent active-state G_q_ and/or G_11_.

As our primary interest is signaling from endosomes, we next asked if we can detect active-state G proteins at endogenous levels on the endosome membrane. To do so, we modified the assay to measure complementation of SmBit-mApple-KB1691 with an endosome-targeted LgBit construct (endofin-LgBit, Fig. 4e). VIP produced a robust endosomal signal, and with slower kinetics than recruitment on the plasma membrane (Fig. 4f). Interestingly, this endosomal signal was largely unaffected by YM (Fig. 4f), in contrast to the plasma membrane signal, which clearly included a non-G_s_, YM-sensitive component (Fig. 4d). These results indicate that the biosensor can indeed detect active-state G_s_ on endosomes, as well as on the plasma membrane, and at native levels. In addition, they suggest that VIPR1 mediates G protein activation in a ‘location biased’ manner, as defined by the production of two distinguishable activation components (G_s_ and non-G_s_) on the plasma membrane but primarily one component (G_s_) on endosomes.

### Active-state Gα_s_ production on endosomes is endocytosis-dependent

We next asked if producing active-state Gα_s_ on endosomes requires the presence of activated GPCRs in the endosome membrane. This is expected based on the present understanding that G protein activation on endosomes requires a second GPCR-G protein coupling reaction that occurs locally on the endosome limiting membrane^4,5^. However, we observed that the GPCR-triggered intracellular redistribution of Gα_s_ to endosomes does not require the presence of receptors on endosomes (Fig. 2, Supplementary Fig. 3). Therefore, we wanted to explicitly test whether or not the production of active-state Gα_s_ on endosomes is endocytosis-dependent, taking advantage of the ability of the KB1691-derived biosensor to detect active-state Gα_s_ on endosomes.

To ask this question, we inhibited VIPR1 internalization using dominant negative (K44E) mutant dynamin^15^. Mutant dynamin reduced the amount of active-conformation VIPR1 at endosomes after VIP application, as detected by a previously described active-state GPCR binder (mini-Gs)^15^, and it increased the amount of active-state VIPR1 at the plasma membrane (Fig. 5a-c), verifying inhibition of receptor endocytosis. Using the KB1691-derived biosensor, we found that mutant dynamin decreased the active-state Gα_s_ signal specifically on the endosome membrane, and to a comparable degree as the decrease in activated receptors in endosomes (Fig. 5d-f, Supplementary Fig. 8). These observations support the conclusion that the production of active-state Gα_s_ on the endosome membrane requires the presence of activated receptors in endosomes, presumably to mediate local GPCR-G protein coupling.

**Figure 5:**
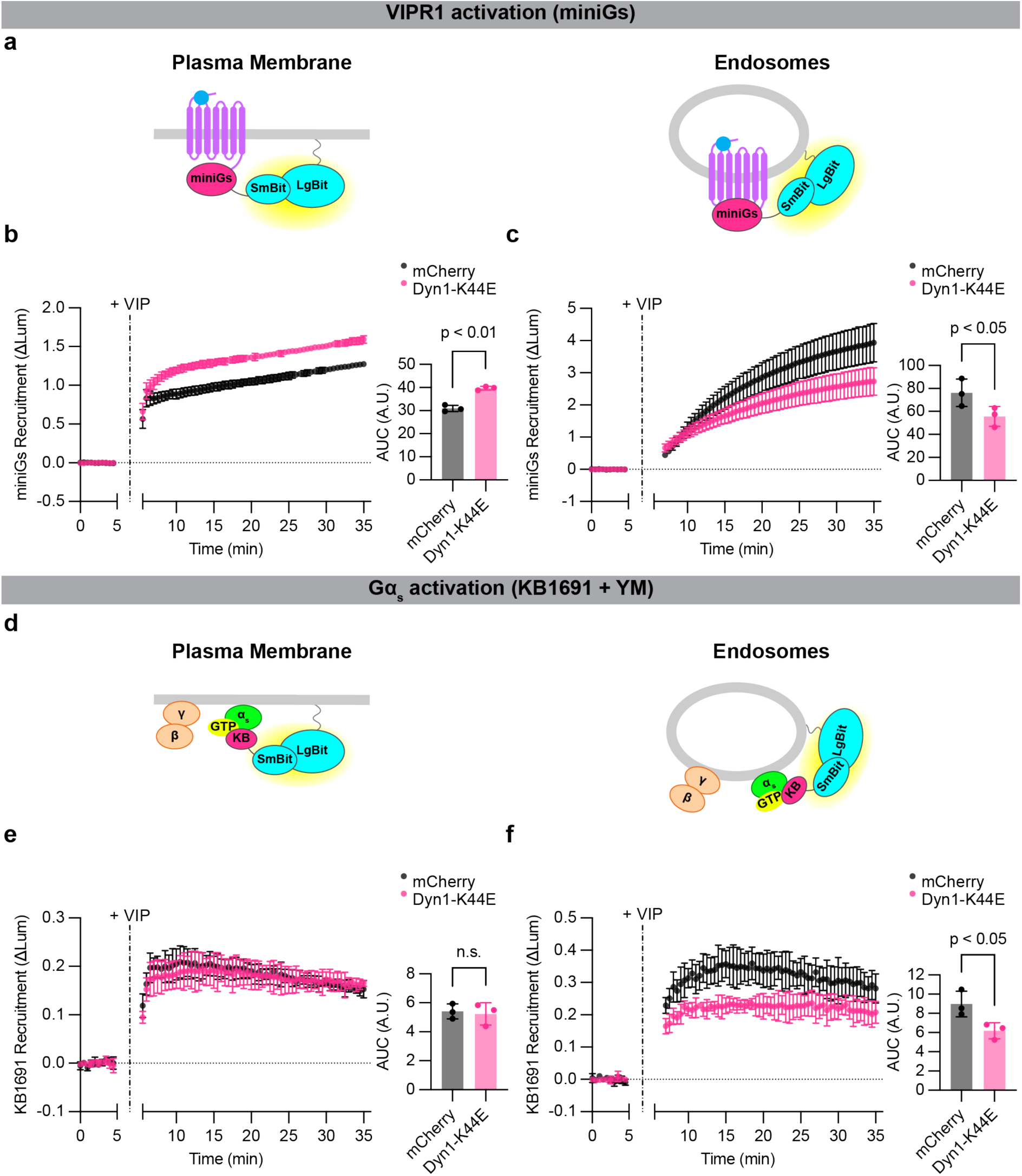
Active-state Gα_s_ production on endosomes depends on VIPR1 endocytosis. **a)** Schematics of mini-Gs (active receptor biosensor) plasma membrane (left) or endosome (right) NanoBit bystander assays. **b,c)** NanoBit bystander assays showing recruitment of mini-Gs to the plasma membrane **(b)** or endosomes **(c)** after activation of Halo-VIPR1 in HEK293 cells transduced with mCherry control or mCherry-Dyn1-K44E BacMam. Left: time course; Right: AUC quantification. VIP (1 µM) was added after 5 minutes. Significance determined by paired two-tailed t-test. **d)** Schematics of KB1691 (active-state Gα_s_ biosensor) plasma membrane (left) or endosome (right) NanoBit bystander assays. **e,f)** Left: NanoBit bystander assay showing Gα_s_-specific recruitment of KB1691 to the plasma membrane **(e)** or endosomes **(f)** after activation of Halo-VIPR1 in HEK293 cells transduced with mCherry control or mCherry-Dyn1-K44E BacMam and pretreated with YM-254890 (1 µM, 30 minutes). Right: AUC of time course. VIP (1 µM) was added after 5 minutes. Data for DMSO-treated control cells are shown in Supplementary Figure 8. Significance was determined by repeated measures 2-way ANOVA with Sidak’s multiple comparisons test with DMSO control data shown in Supplementary Figure 8 (see Supplementary Table 12). Data are shown as mean ± S.D. of 3 independent experiments.

### Detection of VIPR1-mediated coupling to Gα_s_ on endosomes

The results described above indicate that the production of active-state Gα_s_ is endocytosis-dependent, consistent with local coupling on endosomes. We next wanted to investigate whether it is possible to directly detect this proposed coupling reaction with native G proteins. To do so, we adapted the NanoBit assay to detect recruitment of Nb37 rather than KB1691. Nb37 is a nanobody that binds to the N-terminal helical domain of Gα_s_ when destabilized by coupling to receptors or in a nucleotide-free state^36^. This contrasts with KB1691 that binds specifically to the GTP-bound active state^33^, and we focused on Nb37 as a proxy measure of the GPCR-G protein coupling reaction^9^. This approach was shown previously to robustly detect coupling on endosomes in cells overexpressιng both VIPR1 and G_s_^15^, and we found it possible to detect coupling in cells expressing native G proteins using the same experimental design provided that VIPR1 was overexpressed (Fig. 6a-c). VIP produced a robust Nb37 recruitment signal both on the plasma membrane and endosomes, and with the expected sequential kinetics (Fig. 6c). These results, which we interpret as an indication of local GPCR-G protein coupling, were similar to those obtained with the KB1691-derived active-state sensor. Together, these results support the hypothesis that the production of active-state Gα_s_ on endosomes is mediated by local receptor coupling at this location.

**Figure 6:**
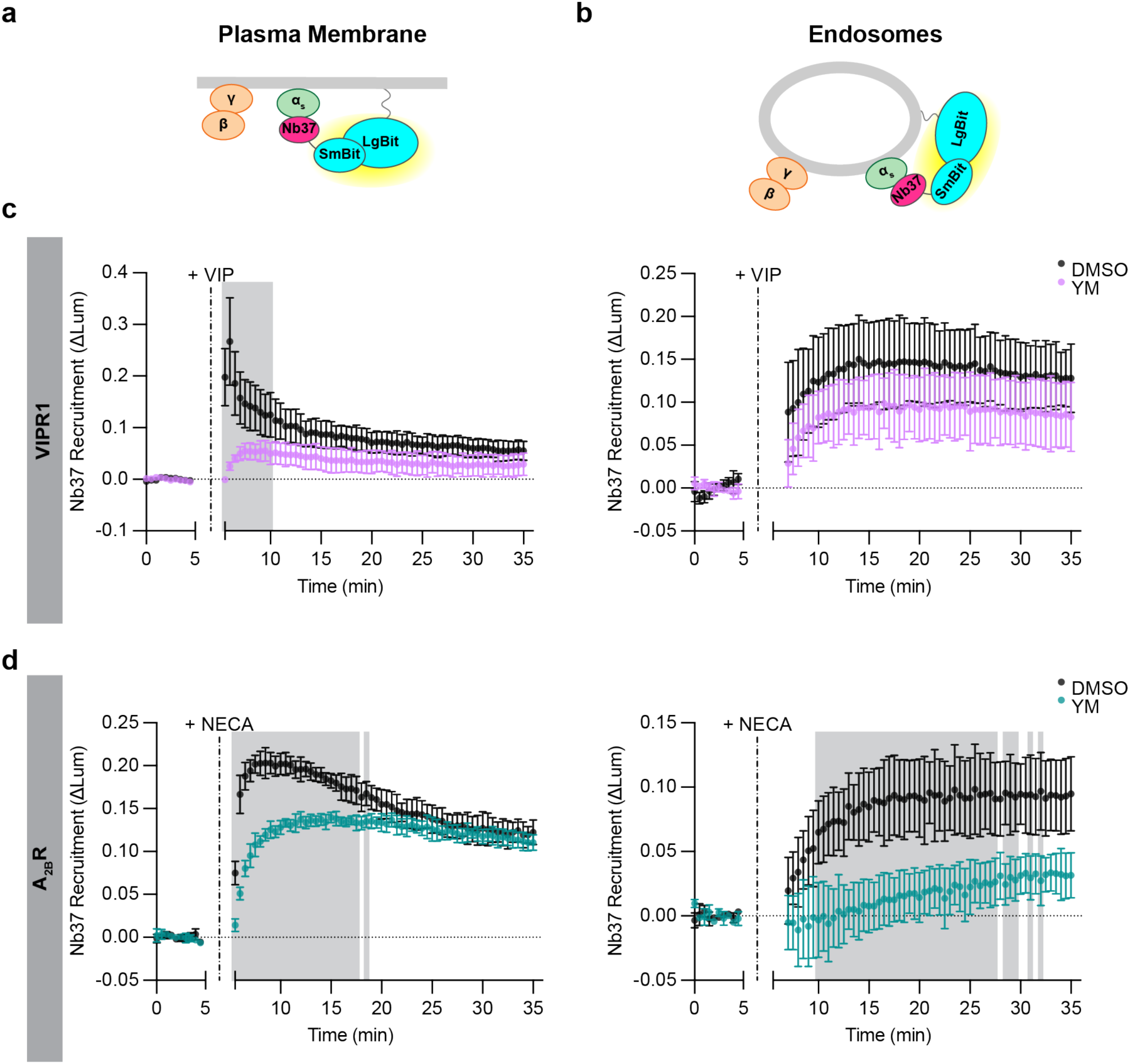
Differences in endosomal G protein activation by VIPR1 and A_2B_R. a,b) Schematics of Nb37 (GPCR-Gα_s_ coupling biosensor) plasma membrane **(a)** and endosome **(b)** NanoBit bystander assays. **c)** NanoBit bystander assays depicting Nb37 recruitment to the PM (left) or endosomes (right) after Halo-VIPR1 activation in HEK293 cells pretreated with either DMSO or YM-254890 (1 µM, 30 minutes). Cells were treated with VIP (1 µM) at 5 minutes. **d)** NanoBit bystander assays depicting Nb37 recruitment to the PM (left) or endosomes (right) after Halo-A_2B_R activation in HEK293 cells pretreated with either DMSO or YM-254890 (1 µM, 30 minutes). Cells were treated with NECA (100 µM) at 5 minutes. Shaded areas in panels c and d represent time points at which the difference between DMSO- and YM-treated cells are statistically significant (p < 0.05, determined by repeated measures 2-way ANOVA with Sidak’s multiple comparisons test, see Supplementary Tables 13 and 14). Data are shown as mean ± S.D. of at least 3 independent experiments.

Because KB1691 detected a non-G_s_ component of the active-state G protein signal after VIPR1 activation, we wanted to also verify specificity of the coupling signal detected by Nb37. We observed a residual signal that remained in Gα_s/olf_ DKO cells, similar to what we observed with KB1691 (Supplementary Fig. 9a). Similar to the non-G_s_ component detected by the KB1691-derived active-state biosensor, the non-G_s_ signal detected by Nb37 recruitment was eliminated in the presence of YM (Supplementary Fig. 9b). Accordingly, we used the same strategy to define G_s_ and non-G_s_ components. Using this approach, we again resolved both G_s_ and non-G_s_ components of VIPR1-mediated coupling on the plasma membrane, but primarily a G_s_ component on endosomes (Fig. 6c). These results provide independent support for the hypothesis that endosomal G_s_ activation is mediated by a local coupling reaction on the endosome limiting membrane, and they verify the existence of distinct G_s_ and non-G_s_ components of G protein activation by VIPR1 on the plasma membrane.

### The location bias of G protein activation on endosomes is GPCR-specific

Our results with VIPR1 support the model that local GPCR-G protein coupling is needed to produce active-state Gα_s_ on endosomes. As an additional approach to test this idea, we took advantage of the fact that the human A_2B_R internalizes weakly after agonist-induced activation relative to VIPR1^31^. If local coupling to activated receptors is required for endosomal G protein activation to occur, we predicted that the A_2B_R would produce a relatively weak G_s_ coupling signal on endosomes. This was indeed the case. A_2B_R activation by NECA clearly produced both G_s_ and non-G_s_ components of Nb37 recruitment at the plasma membrane, but little or no signal representing G_s_ coupling was detected on endosomes (Fig. 6d, Supplementary Fig. 9cd). These results provide additional support for the model that endosomal activation of Gα_s_ requires local receptor-G_s_ coupling.

While A_2B_R activation produced little or no G_s_ coupling signal on endosomes, it did produce a detectable non-G_s_ component (Fig. 6d). This further supports the concept of location bias in G protein coupling on endosomes, relative to the plasma membrane. Remarkably, A_2B_R produced largely the non-G_s_ coupling signal on endosomes, in contrast to VIPR1, which produced largely the G_s_ component, but each GPCR produced both components on the plasma membrane. These results further suggest that distinct GPCR family members are able to differentially bias endosomal G protein activation in a receptor-specific manner (Fig. 7).

**Figure 7:**
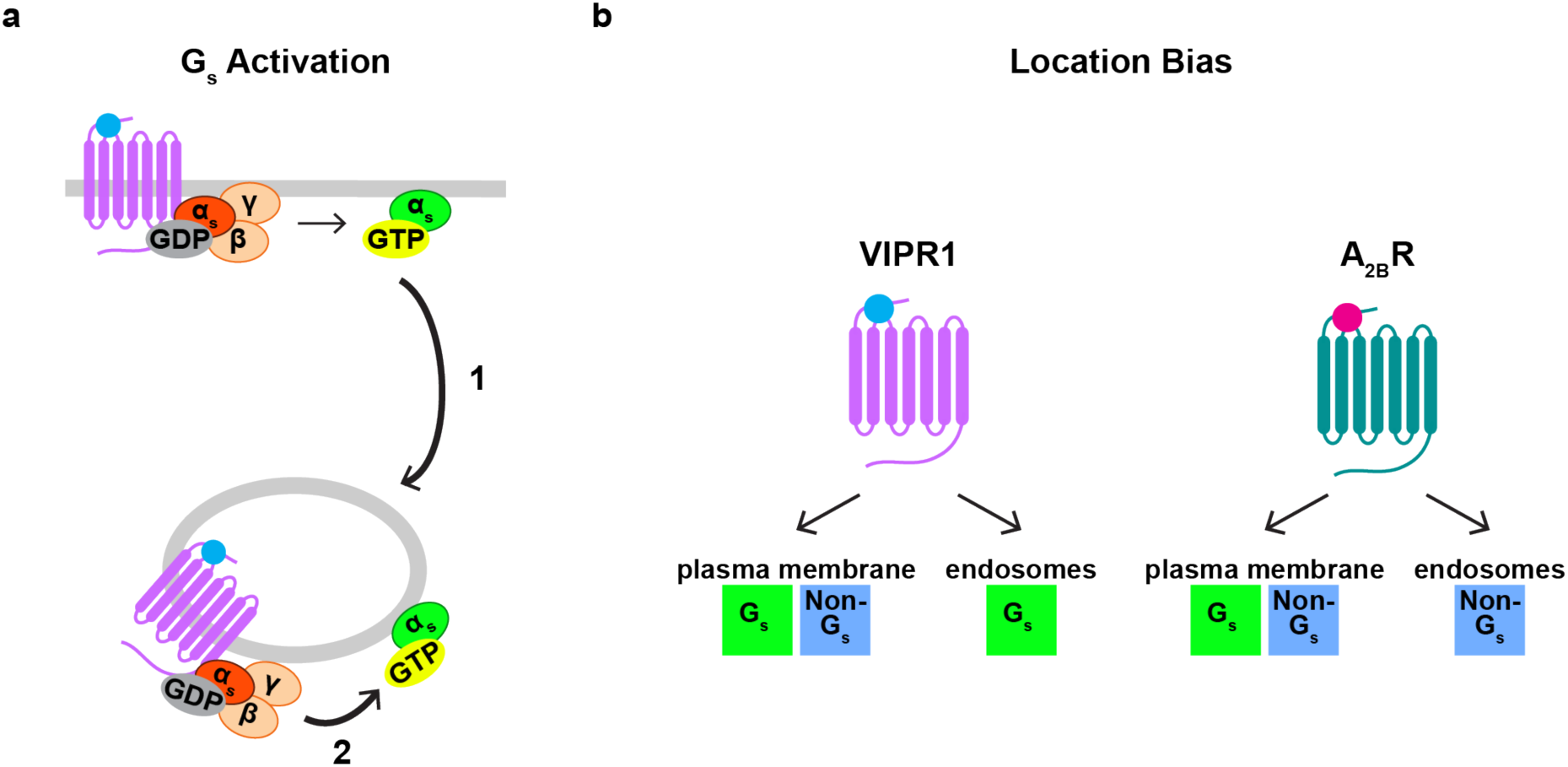
Proposed models of regulation of endosomal G protein activity. **a)** Regulation of endosomal Gα_s_ abundance and activity. Coupling of Gα_s_ to a GPCR at the plasma membrane regulates the abundance of Gα_s_ on endosomes, as it promotes redistribution of Gα_s_ to endosomes independently of GPCR endocytosis (1). A second GPCR-Gα_s_ coupling reaction on endosomes regulates the activity of Gα_s_ on endosomes, which depends on receptor endocytosis (2). **b)** Model of location bias in G protein activation by VIPR1 and A_2B_R. VIPR1 activates both a G_s_ and non-G_s_ component of G protein activity on the plasma membrane but preferentially activates the G_s_ component on endosomes. In contrast, A_2B_R activates both a G_s_ and non-G_s_ component of G protein activity on the plasma membrane but preferentially activates the non-G_s_ component on endosomes.

## Discussion

Many GPCRs are now known to exist in an activated conformation on both endomembranes and the plasma membrane^5^, and there is substantial evidence that they can produce distinct and additional signaling effects from endomembranes^8,31,37–46^. Such signaling requires GPCRs to increase G protein activity on the appropriate endomembrane compartment, but how this is achieved remains unclear. We investigated this question by focusing on how G_s_-coupled receptors control G protein abundance and activity on endosomes.

The ability of G_s_ to redistribute between membranes was proposed more than 40 years ago based on in vitro biochemical reconstitution^47^. Multiple groups have since demonstrated redistribution of Gα_s_ in intact cells and provided insight into its mechanistic basis^19–24^. Here, we began by verifying the present view that 1) G_s_-coupled GPCRs trigger the rapid redistribution of Gα_s_ from the plasma membrane to sample various intracellular membranes, including endosomes, 2) this process is reversible, and 3) it does not require the triggering GPCR to internalize^19,22^. We then build on this understanding by showing that a number of G_s_-coupled GPCRs can trigger this intracellular redistribution process at native receptor expression levels. We next carried out a series of experiments to investigate location-specific regulation of endosomal Gα_s_ activity using conformational biosensors, and we demonstrate that this approach is capable of detecting and localizing active-state Gα_s_ at native G protein levels.

Our results support a simple model in which the abundance and activity of Gα_s_ on endosomes are separately controlled by distinct GPCR-G protein coupling reactions occurring at different subcellular locations–with coupling on the plasma membrane increasing endosomal Gα_s_ abundance and local coupling on endosomes increasing endosomal Gα_s_ activity (Fig. 7a). We detect a small amount of Gα_s_ on endosomes in cells prior to agonist exposure, as noted previously by others^19,27^. Thus, it remains to be determined if the first coupling reaction, which increases Gα_s_ abundance on endosomes, is necessary for signaling from endosomes, or if the basal level of endosomal G_s_ abundance is sufficient. We note that previous studies have come to different conclusions on this question, albeit based on studies of different G protein classes^18,19^.

To our knowledge, the present results are the first to detect and localize active-state Gα_s_ in intact cells expressing only native G proteins. In evaluating the specificity of the biosensors used to achieve this, we were surprised to observe that both biosensors detect a distinct non-G_s_ component, defined by its different kinetics and stimulation in Gα_s/olf_ DKO cells. We currently speculate that this component represents activation of G_q_ and/or G_11_ because it is eliminated by YM, but further study will be necessary to more fully define this component and investigate its potential functional significance. Whereas both VIPR1 and A_2B_R produced both components at the plasma membrane, they selectively produced one component or the other on endosomes in a receptor-specific manner (Fig. 7b). These results support the existence of ‘location bias’ in GPCR-G protein coupling selectivity on endomembranes, consistent with previous evidence for such bias both through assays of functional signaling^7,48^ and recruitment of different biosensors of GPCR activation^49–51^. Further, they suggest that this location bias is GPCR-specific.

It is also remarkable that A_2B_R produces a detectable non-G_s_ coupling signal on endosomes, despite this GPCR internalizing only weakly when compared to VIPR1^31^. This observation is consistent with a previous study indicating that G_q/11_ activation on endosomes can occur even when the concentration of activated receptors in endosomes is reduced to a low level by endocytic inhibition^18^. We also note emerging evidence for the existence of a second cellular mechanism of G_i/o_ activation on endosomes that does not depend on the activating GPCRs being present in endosomes^52^.

In closing, the present results add to the currently expanding mechanistic framework of spatiotemporal GPCR signaling through heterotrimeric G protein. They also raise new questions that may help to guide further elucidation of this process and enable its future therapeutic manipulation.

## Methods

### Cell culture and transfections

HEK293 cells were purchased from ATCC (CRL-1573) and cultured in DMEM (Gibco 11965-092) and 10 % FBS (R&D Systems, S12495) at 37 °C and 5 % CO_2_ in a humid environment. All cell lines used in this study were generated from HEK293 cells, with the exception of HEK293A *GNAS/GNAL* double knockout^53^ cells used in Supplementary Figure 5. Polyclonal cells stably expressing Flag-β2AR^31^ were cultured in 500 μg/mL geneticin (Gibco 10131027). All cell lines were routinely screened for mycoplasma contamination (MycoAlert, Lonza LT07-318). Cells were transfected using Lipofectamine 2000 (Thermo Fisher 11668019) according to the manufacturer’s protocol. For experiments in Figure 5 and Supplementary Figure 8, cells were transfected with appropriate receptor and nanobit pairs and transduced after four hours with either mCherry or mCherry-Dynamin-1 K44E BacMam diluted in culture media.

### DNA constructs and molecular cloning

All DNA constructs used in this study are listed in Supplementary Table 1. Novel constructs were constructed by standard InFusion (Takara Bio) or KLD (NEB) cloning techniques, following the manufacturers’ protocols, and sequences were confirmed by Sanger sequencing. mCherry and mCherry-Dynamin-1 K44E BacMam produced from pCMV-Dest (Thermo Fisher A24223) used in Figure 5 and Supplementary Figure 8 were produced according to the manufacturer’s protocol.

### Generation of CRISPR KO cell lines

Single guide RNAs (sgRNAs) were designed with the Synthego CRISPR design tool (Supplementary Table 2). To generate ribonucleoproteins (RNPs), 3 μL of 53.3 μM sgRNA (Synthego) were mixed with 2 μL of 40 μM Cas9 (UC Berkeley Macrolab) and incubated at room temperature for 10 minutes. Cells (2.0 x 10^5^) were prepared for electroporation with RNPs with the SF Cell Line 4D Nucleofector kit (Lonza) following the manufacturer’s protocol and electroporated in a 4D Nucleofector (Lonza) using program CM-130. After electroporation, monoclonal cell lines were established using standard techniques and genetic modifications were verified using either Sanger sequencing or next-generation sequencing (Amplicon-EZ, Azenta Life Sciences, see Supplementary Table 3 for NGS primers). For novel Gα_s/olf_ DKO cells, modifications were done sequentially.

### Generation of EGFP-Gα_s_ KO cell lines

Gα_s_ KO1 and KO2 cells were transfected with EGFP-Gα_s_ and selected with 500 μg/mL geneticin. After selection, cells expressing EGFP-Gα_s_ were sorted into a polyclonal population using a FACSAria Fusion Flow Cytometer (BD Biosciences, KO1) or FACSAria III flow cytometer (BD Biosciences, KO2). Cells were cultured under continued selection.

### Live cell confocal microscopy

Confocal imaging was performed using a fully automated Nikon Ti inverted microscope equipped with a CSU-22 spinning disk (Yokogawa), piezo stage (Mad City Labs), 4-line Coherent OBIS laser launch (100 mW at 405, 488, 561, and 640 nM), a quad dichroic 405/491/561/640 (Yokogawa), and corresponding emission filters ET460/50m, ET525/50m, ET610/60m, ET700/75m in a filter wheel controlled by a Lambda 10-3B (Sutter) for channels DAPI/GFP/RFP/Cy5, respectively. Images were captured using an Apo TIRF 100x/1.49 oil objective lens (Nikon) and a Photometrics Evolve Delta EMCCD Camera (154 nm/pixel) controlled with Nikon NIS Elements HC v.5.21.03 software.

For live imaging, cells grown in either 6-well plates or 6 cm dishes were transfected 48 hours before imaging and plated into 35 mm glass bottom microscopy dishes (Cellvis D35-20-1.5-N) coated with 0.001 % (w/v) poly-L-lysine (Millipore Sigma P8920) 24 hours after transfection. Receptors were surface labelled with either monoclonal anti-Flag M1 antibody (Millipore Sigma F3040) labelled with Alexa Fluor 647 (Thermo Fisher A20186) or 200 nM JF_635_i-HTL^54^ for 10 minutes at 37 °C and 5 % CO_2_. After labelling, cells were washed three times and imaged in imaging media (DMEM (no phenol red, Gibco 31053-028) supplemented with 30 mM HEPES pH 7.4) in a temperature- and humidity-controlled chamber at 37 °C (OkoLab). Time-lapse images in Figure 1, Supplementary Figure 1, Supplementary Figure 3, and Supplementary Figure 4 were acquired by imaging cells at 20 second intervals for 30 minutes, and agonist (specified in figure legends) was added after 5 minutes. In Supplementary Figure 3a, cells were treated with either Dyngo4a (30 µM, Abcam ab120689) or DMSO (0.1 %) for 25 minutes prior to imaging, Alexa Fluor 647 coupled anti-Flag M1 antibody was added for the last 10 minutes of pretreatment, and cells were washed and imaged in imaging media containing either Dyngo4a or DMSO. For time lapse images in Figure 2 and Supplementary Figure 2, cells were imaged at 20 second intervals for 60 minutes with 100 nM Iso added at 5 minutes and 10 μΜ Alp added at 15 minutes.

Images were processed for presentation in Fiji v2.14^55^. Pearson correlation analysis was performed using Cell Profiler v4.2.6^56^; briefly, cells were segmented based on the green channel for each frame and Pearson correlation was calculated between channels at each time point.

### Fixed imaging

Cells were transfected in 6 well plates 48 hours before fixation. After 24 hours, cells were split onto coverslips coated with 0.001 % poly-L-lysine (Millipore Sigma P8920) in 12 well plates and grown for an additional 24 hours. Cells were then surface labelled with monoclonal anti-Flag M1 antibody (Millipore Sigma F3040) labelled with Alexa Fluor 647 (Thermo Fisher A20186) for 10 minutes at 37 °C and 5 % CO_2_. After labelling, cells were washed two times with full media (DMEM + 10 % FBS) and treated with either vehicle or Iso (1 µM) for an additional 15 minutes at 37 °C and 5 % CO_2_. Cells were then placed on ice, washed 1x with DPBS, and fixed at room temperature for 10 minutes in 3.7 % formaldehyde in modified BRB80 (80 mM PIPES pH 6.8, 1 mM MgCl_2_, 1 mM CaCl_2_). After fixation, cells were washed 3 times with DPBS, incubated in TBS for 20 minutes, and washed an additional 3 times with DPBS. Dapi (1:5000) was included in the final wash. Cells were then mounted in ProLong Gold Antifade mounting medium and left to dry overnight in the dark.

Slides were imaged using the confocal microscope described above using a Plan Apo VC 60x/1.4 oil objective lens. Images were processed for presentation using Fiji v2.14^55^.

### TIRF microscopy

For live cell TIRF microscopy, cells were imaged using the same methods as for live cell confocal microscopy. Images were acquired on a fully automated inverted Nikon Ti-E microscope controlled by Nikon NIS-Elements software (5.20.00 build 1423), a Nikon motorized stage equipped with a TIRF module with STORM lens (Nikon), Agilent MLC400 (405nm, 488nm, 561nm, 647nm) light source with NIDAQ interface (v18.00), and corresponding emission filters ET455/50m, ET525/50m, ET600/60m, ET705/72m in a filter wheel controlled by a Lambda 10-3B (Sutter) for channels DAPI/GFP/RFP/Cy5, respectively. Images were captured using an Apo TIRF 100x/1.49 objective (Nikon) with an Andor DU897 EMCCD camera and an OkoLab temperature controlled live stage insert. Time lapse images were acquired at 10 second intervals for 25 minutes at 37 °C with agonist added at 5 minutes.

Images were analyzed and processed for presentation using Fiji v2.14^55^. To quantify relative changes in surface fluorescence (F/F_0_), cells were manually segmented by drawing a region of interest (ROI) around the cell surface. The mean, background subtracted, fluorescence intensity (F) was measured at each time point and normalized to the average mean fluorescence intensity before agonist treatment (F_0_). To calculate vehicle subtracted F/F_0_ values, the F/F_0_ of vehicle control replicates were averaged at each time point and subtracted from the F/F_0_ values at each time point for individual movies. For presentation in Figure 3 and Supplementary Figure 7, images were scaled to the pre-agonist time point and pseudo-colored using the viridis colormap in Fiji to visualize relative changes in fluorescence.

### NanoBit luciferase complementation assays

Cells were grown in 6-well plates or 6 cm dishes and transfected with both receptor constructs (Flag-β2AR, Halo-VIPR1, or Halo-A_2B_R) and the appropriate LgBit- and SmBit-tagged constructs (in Nb37 assays shown in Figure 6 and Supplementary Figure 9, the SmBit(101) tag was used, while the SmBit(114) tag was used in all other assays^57^). After 24 hours, cells were washed, lifted, spun at 500 x g for 3 minutes and resuspended in assay buffer (20 mM HEPES pH 7.4, 135 mM NaCl, 5 mM KCL, 0.4 mM MgCl_2_, 1.8 mM CaCl_2_) with 5 µM coelenterazine-H (Research Product International C61500). Cells were plated (100 µL) into untreated white 96-well plates (Corning 3912) and incubated at 37 °C and 5 % CO_2_ for either 10 minutes or 30 minutes (for cells pretreated with DMSO (0.1 %) or YM-254890 (1 μM, Cayman Chemical 29735 or Tocris 7352), as indicated in figure legends) before reading. Luminescence was measured on either a Synergy H4 (BioTek, for data in Figure 2 and Supplementary Figure 3) or Spark (Tecan, for all other data) plate reader. For assays in Figure 2, luminescence was read every 1 minute for a 5-minute baseline, after which either vehicle or Iso (100 nM) was added and luminescence read for 10 minutes, followed by vehicle or Alp (10 µM) addition and additional luminescence reading for 30 minutes. For assays in Supplementary Figure 3, luminescence was read every 1 minute for a 5-minute baseline, after which either vehicle or Iso (1 µM) was added and luminescence read for an additional 30 minutes. For all other assays, luminescence was read every 30 seconds for a 5-minute baseline, after which vehicle or agonist (noted in figure legends) was added and luminescence measured for an additional 30 minutes. For cells pretreated with DMSO or YM-254890, cells were kept in continual treatment for the duration of the assay.

To analyze data, the change in normalized luminescence was calculated by normalizing each well to its average baseline luminescence. Then, the average change of luminescence of vehicle-treated wells was subtracted from the average change in luminescence of agonist-treated cells. Data are represented as the vehicle subtracted change in luminescence of agonist-treated cells (ΔLum = Lum_agonist_ - Lum_vehicle_).

### cADDis cAMP assays

Intracellular cAMP levels were measured using either Green Up cADDis cAMP biosensor (Montana Molecular U02006, Supplementary Figure 5) or Red Up cADDis cAMP biosensor (Montana Molecular U0200R, Supplementary Figure 6) following the manufacturer’s protocol. Briefly, 50,000 cells were plated into TC-treated black 96-well plates (Corning 3340) coated with 0.001 % (w/v) poly-L-lysine (Millipore Sigma P8920) and transduced with cADDis BacMam. After 24 hours, cells were washed with assay buffer (20 mM HEPES pH 7.4, 135 mM NaCl, 5 mM KCL, 0.4 mM MgCl_2_, 1.8 mM CaCl_2_) twice and incubated at 37 °C in a temperature controlled plate reader (Tecan Spark for Green cADDis assays or BioTek Synergy H4 for Red cADDis assays). Baseline fluorescence was read with an excitation wavelength at 500 nm (Green cADDis) or 558 nm (Red cADDis) and emission wavelength at 530 nm (Green cADDis) or 603 nm (Red cADDis) for 5 minutes every 30 seconds, after which agonist (noted in figure legends) was added and fluorescence read for an additional 30 minutes. To calculate the change in intracellular cAMP (ΔF/F_0_), the average baseline fluorescence for each well was calculated (F_0_) and the change in fluorescence for each well (ΔF = F - F_0_) was normalized to F_0_.

### Immunoblotting

Cells were lysed in RIPA buffer (50 mM Tris pH 7.4, 150 mM NaCl, 1 % Triton X-100, 0.5 % sodium deoxycholate, 0.1 % SDS) supplemented with Roche cOmplete EDTA-free protease inhibitor tablets (Roche 04693159001) and lysate was boiled at 95 °C for 5 minutes in NuPage LDS Sample Buffer (Thermo Fisher, NP0007) and 20 mM DTT. For Flag-β2AR blots in Supplementary Figure 3, lysate was incubated with LDS Sample Buffer and 20 mM DTT for 1 hour at room temperature instead of boiled at 95 °C. SDS-PAGE and western blots were performed using standard techniques with a polyclonal rabbit anti-Gα_s/olf_ antibody (LS-Bio LS-B4790, 1:1000, blocked in Tris-buffered saline, 5 % milk, 0.1 % Tween-20), a monoclonal rabbit anti-GAPDH antibody (D16H11, Cell Signaling Technology 5174S, 1:1000, blocked in LICOR Intercept (TBS) blocking buffer (Licor 927-60001), a monoclonal rabbit βARR1/2 antibody (D24H9, Cell Signaling Technology 4674, 1:1000, blocked in LICOR Intercept (TBS) blocking buffer), or a monoclonal mouse anti-Flag M1 antibody (Millipore Sigma F3040, 1:1000, blocked in LICOR Intercept (TBS) blocking buffer), followed by IRDye 800- or 680-linked anti-mouse or anti-rabbit IgG secondary antibodies (LI-COR Biosciences). Blots were imaged using an Odyssey Imager (v.2.0.3, LI-COR Biosciences) and quantified using Fiji (v2.14).

### Statistical analysis and reproducibility

Microscopy quantification data are presented as mean ± S.E.M. of individual dishes from at least 3 independent experiments (with the exception of Figure 3b, which are mean ± S.E.M. of individual dishes from at least 2 independent experiments), while cAMP and NanoBit data are presented as mean ± S.D. from at least 3 independent experiments. Each biological replicate in cAMP and NanoBit assays represents the average of at least 2 technical replicates. All images are representative of at least three biologically independent experiments, with the following exceptions: images in Figure 3a, which are representative of at least two independent experiments, and images in Supplementary Figure 2b, Supplementary Figure 4, and Supplementary Figure 6, which are representative of two independent experiments. Statistical tests and area under the curve calculations were performed using GraphPad Prism (v.9 and v.10).

## Data availability

Data and materials are available upon request.

## Acknowledgements

We thank Nicole Fisher for sharing plasmids; Asuke Inoue and Luke Lavis for sharing reagents and advice; Aashish Manglik for sharing equipment; and Natalia Jura, Roshanak Irannejad, Barbara Panning, Rita Fagan, Nicole Fisher, and the rest of the von Zastrow lab for helpful discussion and feedback on the manuscript. Imaging data for this study were acquired at the UCSF Center for Advanced Light Microscopy (Nico Stuurman, Kari Herrington, Micaela Lasser, DeLaine Larsen, and SoYeon Kim). The UCSF Helen Diller Family Comprehensive Cancer Center Laboratory for Cell Analysis (Sarah Elmes, supported by the NIH under award P30CA082103) assisted with cell line generation. This work was supported by the NIH under awards R01DA010711 and R01DA012864 (to M.v.Z.). B.W. was supported by T32GM007810.

## Author contributions

B.W. and M.v.Z. conceptualized the study. B.W., E.E.B., and M.v.Z. designed experiments.

B.W. and E.E.B. performed experiments and analyzed data. B.W. and M.v.Z. wrote the manuscript with input from all authors.

## Competing interests

The authors declare no competing interests.

Correspondence and request for materials should be made to Mark von Zastrow.

