## Supplementary Figures for "Conformational biosensors delineate endosomal G protein regulation by GPCRs"

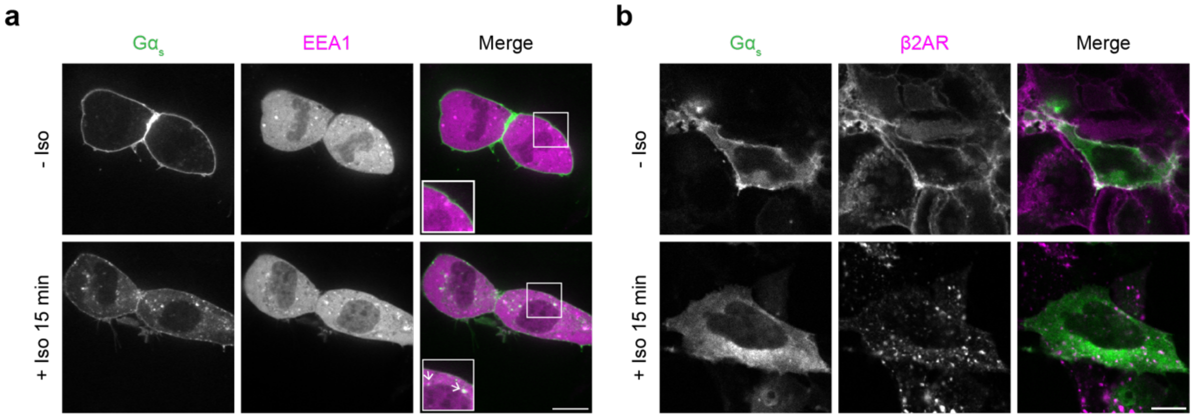


**Supplementary Figure 1: $G\alpha_s$ localizes to endosomes after receptor activation but appears more diffuse after fixation. a)** Representative stills from time-lapse confocal microscopy of live HEK293 cells stably expressing Flag-β2AR and transiently expressing both EGFP-$G\alpha_s$ and the endosomal marker mApple-EEA1 before or after 15 minutes of Iso (1 μM) treatment. **b)** Representative images of fixed HEK293 cells stably expressing Flag-β2AR and transiently expressing EGFP-$G\alpha_s$ before or after 15 minutes of Iso (1 μM) treatment. Prior to imaging (a) or drug treatment and fixation (b), cells were treated for 10 minutes with an anti-Flag antibody coupled to Alexa Fluor 647 to label surface Flag-β2AR. Cells were co-transfected with myc-$G\beta_1$ and untagged $G\gamma_2$. Images are representative of at least 3 independent experiments, scale bars are 10 μm, and insets in panel a are 1.5x zoom on indicated regions. Arrows indicate examples of colocalization.

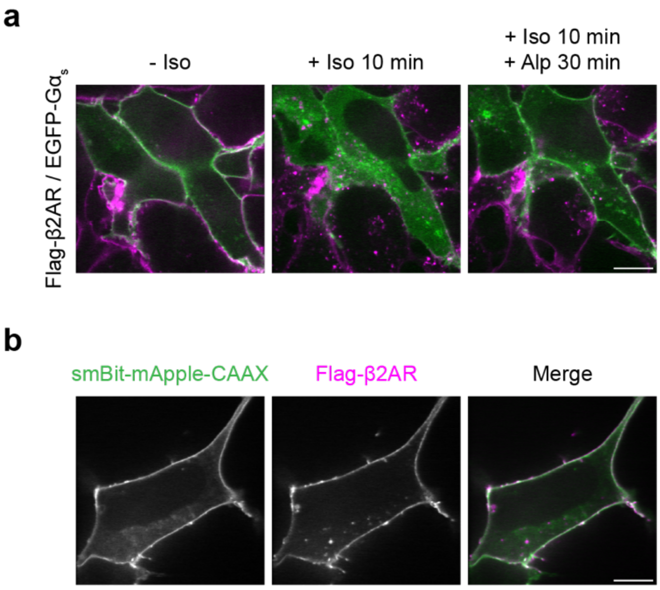


**Supplementary Figure 2: G$\alpha_s$ returns to the plasma membrane after receptor inactivation. a)** Representative stills from time-lapse confocal imaging of live HEK293 cells stably expressing Flag-β2AR and transfected with EGFP-G$\alpha_s$, myc-G$\beta_1$, and untagged G$\gamma_2$ either before drug treatment, after 10 minutes of Iso (100 nM) treatment, or after 10 minutes of Iso followed by 30 minutes of Alprenolol (Alp, 10 μM) treatment. Images are merged G$\alpha_s$ / β2AR channels of images in Figure 2a. **b)** Representative confocal images of HEK293 cells transfected with smBit-mApple-CAAX and Flag-β2AR. Prior to imaging, cells were treated for 10 minutes with an anti-Flag antibody coupled to Alexa Fluor 647 to label surface Flag-β2AR. Images are representative of 4 (a) or 2 (b) independent experiments. Scale bars = 10 μm.

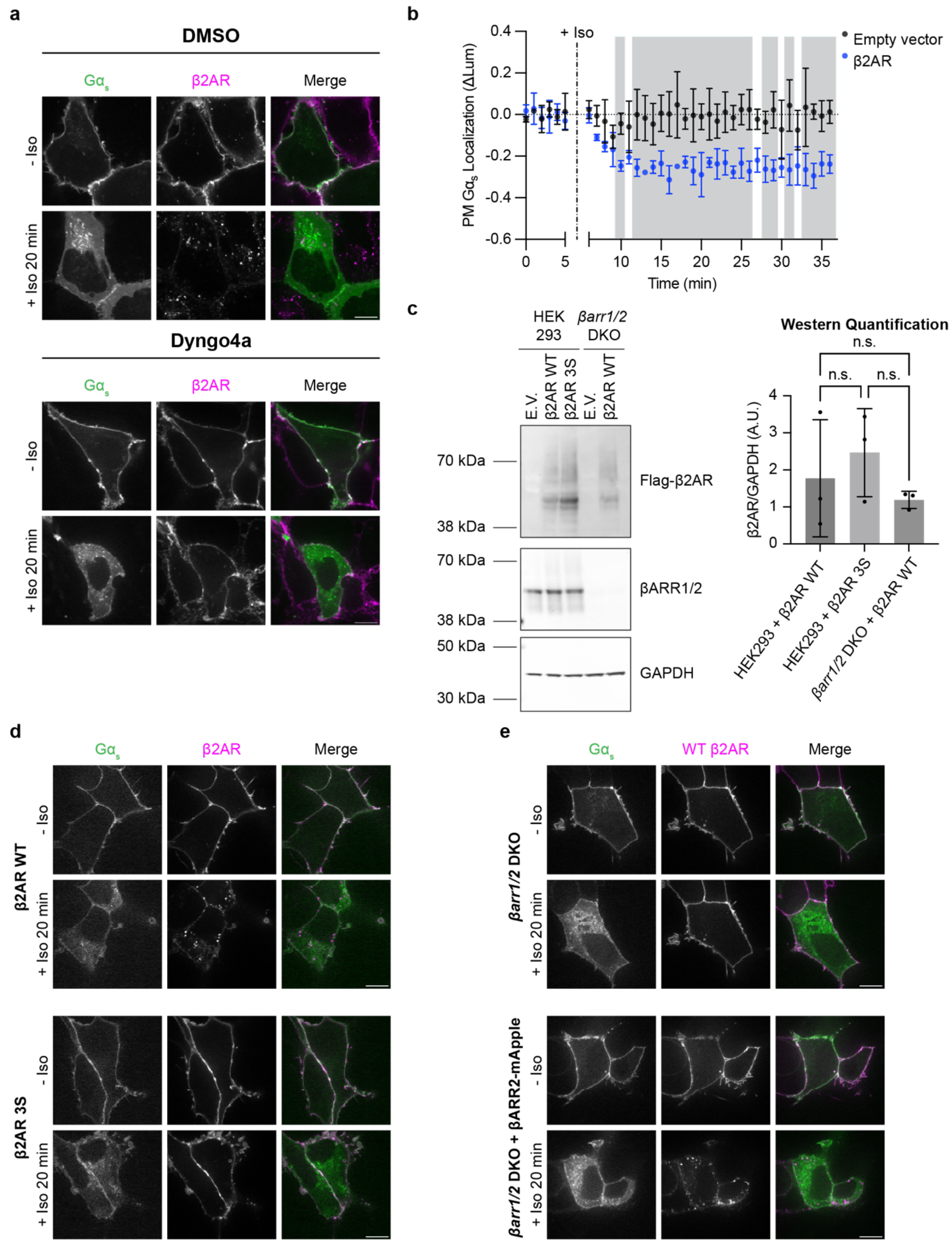


**Supplementary Figure 3: Gα$_s$ redistribution is independent of receptor internalization.** **a)** Representative stills from time-lapse confocal microscopy of live HEK293 cells stably expressing Flag-β2AR and transiently expressing EGFP-Gα$_s$ before and after 20 minutes of Iso (1 μM) treatment. Cells were either pretreated with DMSO (top, 0.1 %) or Dyngo4a (bottom, 30 μM) for 25 minutes prior to imaging, and cells were incubated for the final 10 minutes of compound pretreatment with an anti-Flag antibody coupled to Alexa Fluor 647 to label surface Flag-β2AR. Iso was added after 5 minutes of imaging. Scale bar = 10 μm. **b)** NanoBit bystander assay showing plasma membrane localization of Gα$_s$ in HEK293 cells expressing WT Flag-β2AR (blue) or empty vector (black). Iso (1 μM) was added at 5 minutes. Shaded areas represent time points at which the difference between β2AR and empty vector were significant ($p < 0.5$, determined by repeated measures ANOVA with Sidak's multiple comparisons test, see Supplementary Table 7). **c)** Left: representative western blots of HEK293 cells expressing Flag-β2AR WT or Flag-β2AR 3S, or *βarr1/2* DKO cells expressing WT β2AR. Right: Densitometry analysis of western blots. Significance was determined by ordinary one-way ANOVA followed by Tukey's multiple comparisons test. **d)** Representative confocal images of HEK293 cells transiently coexpressing either WT Flag-β2AR (top) or Flag-β2AR-3S (bottom) and EGFP-G$\alpha_s$ before or after 20 minutes of Iso (100 nM) treatment. Cells were co-transfected with untagged G$\beta_1$ and mApple-G$\gamma_2$. **e)** Representative confocal images of *βarr1/2* DKO HEK293 cells transiently coexpressing WT Flag-β2AR, EGFP-G$\alpha_s$, and either mApple control (top) or βARR2-mApple (bottom). Cells were co-transfected with untagged G$\beta_1$ and myc-G$\gamma_2$ before or after 20 minutes of Iso (100 nM) treatment. In panels d and e, cells were treated prior to imaging for 10 minutes with an anti-Flag antibody coupled to Alexa Fluor 647 to label surface Flag-β2AR. Images are representative of at least 3 independent experiments, and scale bars are 10 μm. Data are represented as mean ± S.D. of 3 independent experiments.

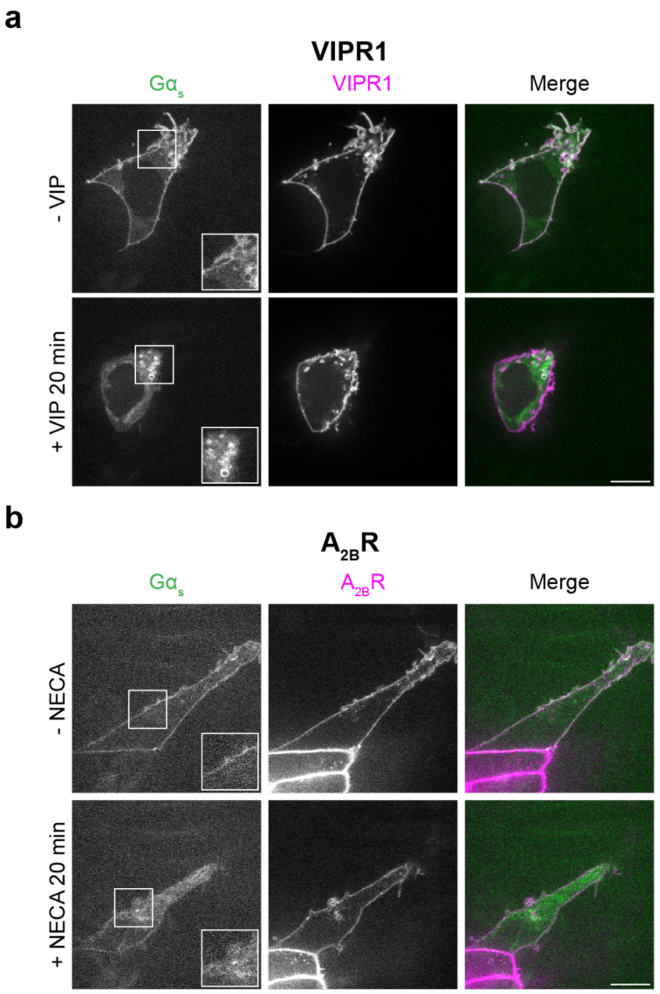


**Supplementary Figure 4: Halo-VIPR1 and Halo-$A_{2B}$R trigger redistribution of G$\alpha_s$ to internal compartments. a,b)** Representative stills from time-lapse confocal microscopy of live HEK293 cells expressing EGFP-G$\alpha_s$ and either Halo-VIPR1 **(a)** or Halo-$A_{2B}$R **(b)** before and after 20 minutes of VIP (1 μM, a) or NECA (100 μM, b) treatment. Insets (1.5x) show examples of internal punctate localization of EGFP-G$\alpha_s$ after receptor activation. Prior to imaging, cells were treated for 10 minutes with 200 nM $JF_{635}$i-HTL to label surface receptors. Images are representative of two independent experiments. Scale bars = 10 μm.

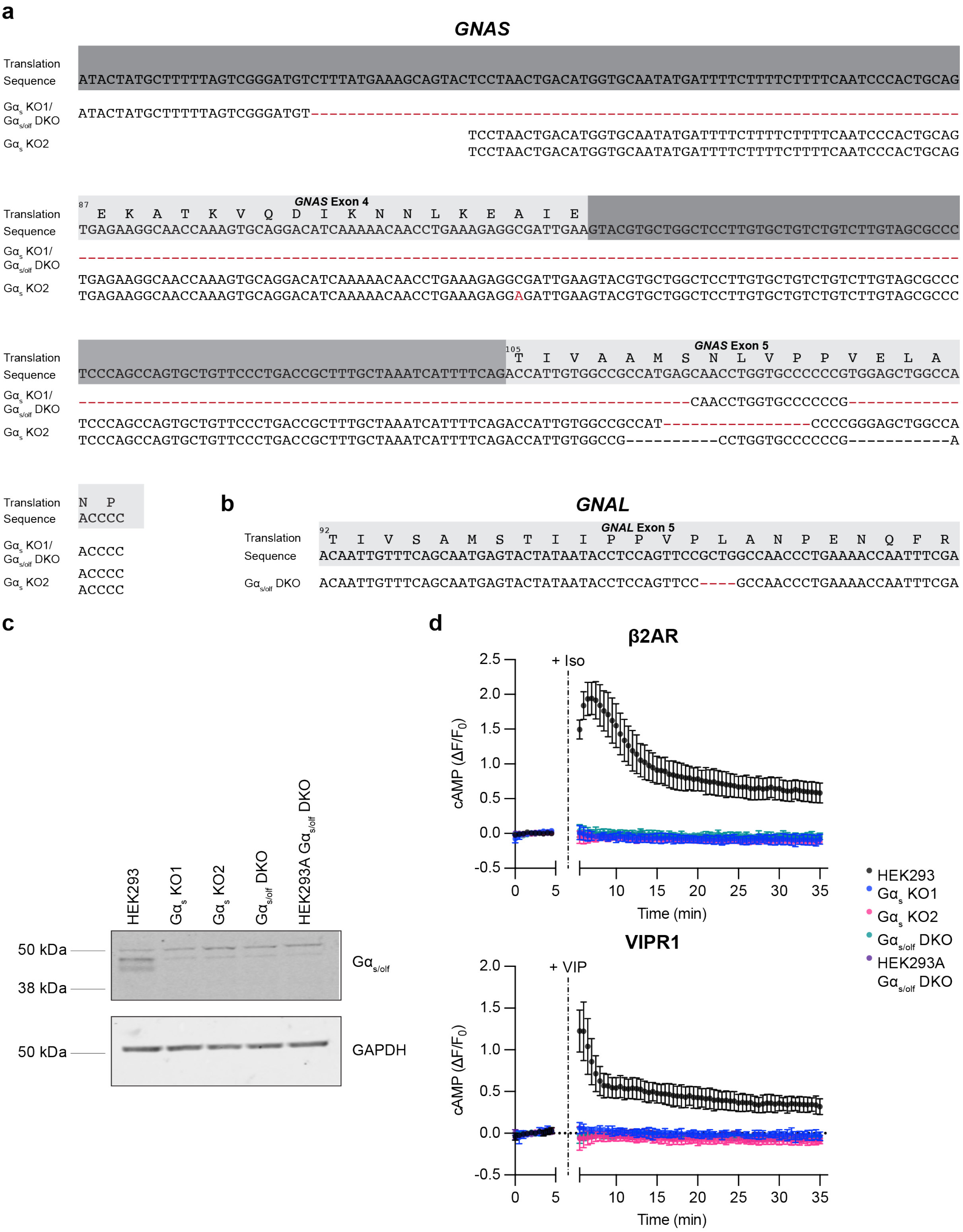


**Supplementary Figure 5: Characterization of Gα$_s$ KO and Gα$_{s/olf}$ DKO cells used in this study. a)** Sequence alignments of *GNAS* locus (encoding Gα$_s$) demonstrating genetic modifications of novel monoclonal Gα$_s$ KO and Gα$_{s/olf}$ DKO cell lines generated in this study. **b)** Sequence alignments of *GNAL* locus (encoding Gα$_{olf}$) demonstrating genetic modifications of the novel monoclonal Gα$_{s/olf}$ cell line generated in this study. **c)** Gα$_{s/olf}$ western blot of Gα$_s$ KO and Gα$_{s/olf}$ DKO cells confirming knockout of Gα$_{s/olf}$ protein. HEK293A Gα$_{s/olf}$ cells[53] were included as a positive control for non-specific antibody binding. Blot is representative of 3 independent experiments. **d)** Green cADDis cAMP assays of HEK293 parental cells and monoclonal G$\alpha_s$ and G$\alpha_{s/olf}$ cells after endogenous β2AR (top) or VIPR1 (bottom) activation. Cells were treated with Iso (100 nM) or VIP (1 μM) at 5 minutes. HEK293A G$\alpha_{s/olf}$ cells[53] were included as a positive control for loss of cAMP response. Data are presented as mean ± S.D. of at least 3 independent experiments.

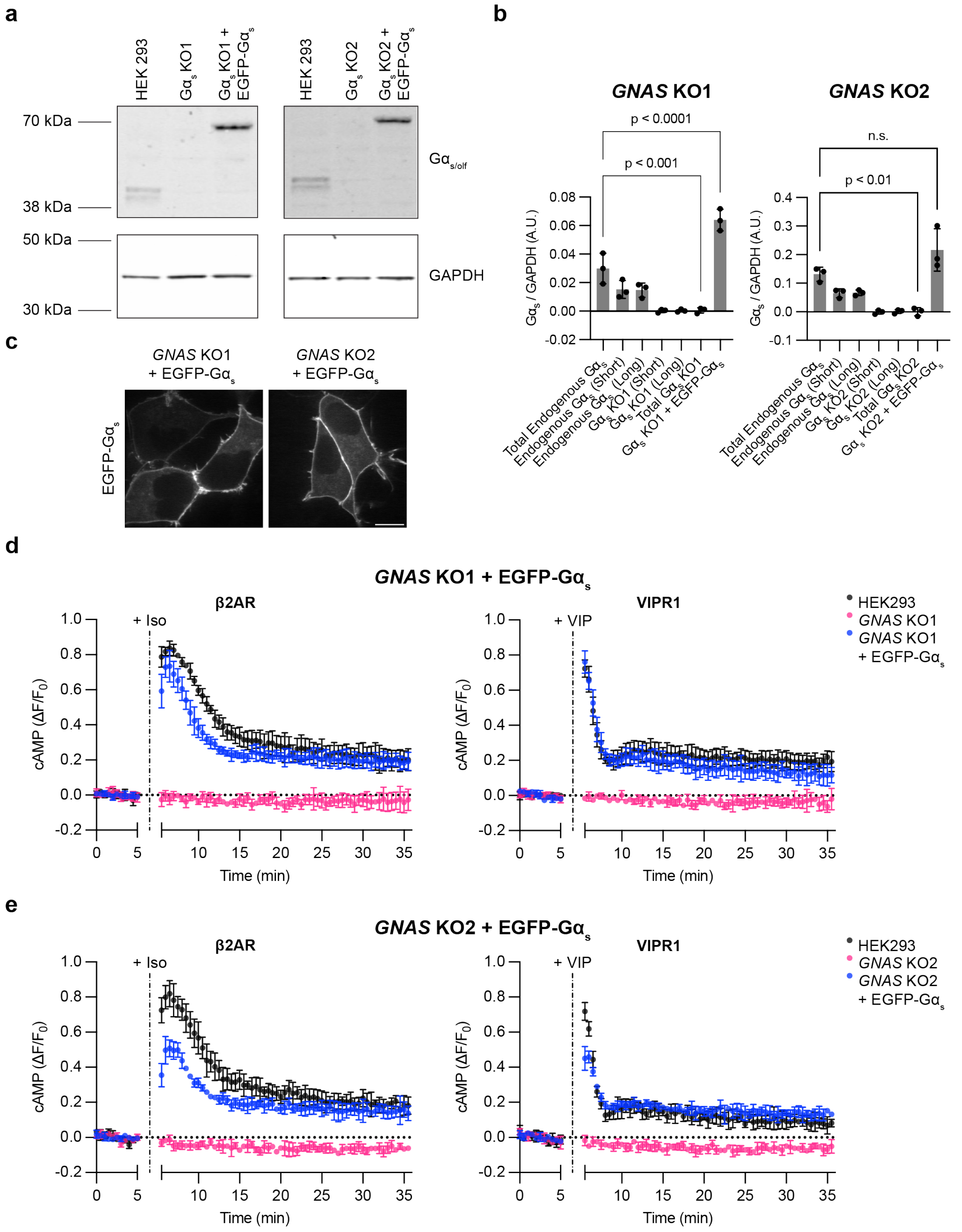


**Supplementary Figure 6: Characterization of Gαs KO + EGFP-Gαs rescue cells used in this study. a)** Representative western blots of parental HEK293, Gαs KO1 cells, and Gαs KO1 + EGFP-Gαs rescue cells (left) or HEK293 cells, Gαs KO2 cells, and Gαs KO2 + EGFP-Gαs rescue cells (right). **b)** Densitometry analysis of western blots from a). Data are shown as mean ± S.D. of 3 independent experiments. Significance was determined by one-way ANOVA followed by Tukey's multiple comparisons test. **c)** Confocal micrographs of live Gαs KO1 or KO2 + EGFP-Gαs rescue cells demonstrating plasma membrane localization of

EGFP-G$\alpha_s$. Images are representative of 2 independent experiments. **d)** Red cADDis cAMP assays of HEK293 parental cells, G$\alpha_s$ KO1, and G$\alpha_s$ KO1 + EGFP-G$\alpha_s$ rescue cells after activation of endogenous β2AR (left) or VIPR1 (right). **e)** Red cADDis cAMP assays of HEK293 parental cells, G$\alpha_s$ KO2, and G$\alpha_s$ KO2 + EGFP-G$\alpha_s$ rescue cells after activation of endogenous β2AR (left) or VIPR1 (right). For data in d) and e), Iso (100 nM) or VIP (1 μM) were added at 5 minutes, and data are shown as mean ± S.D. of at least 3 independent experiments.

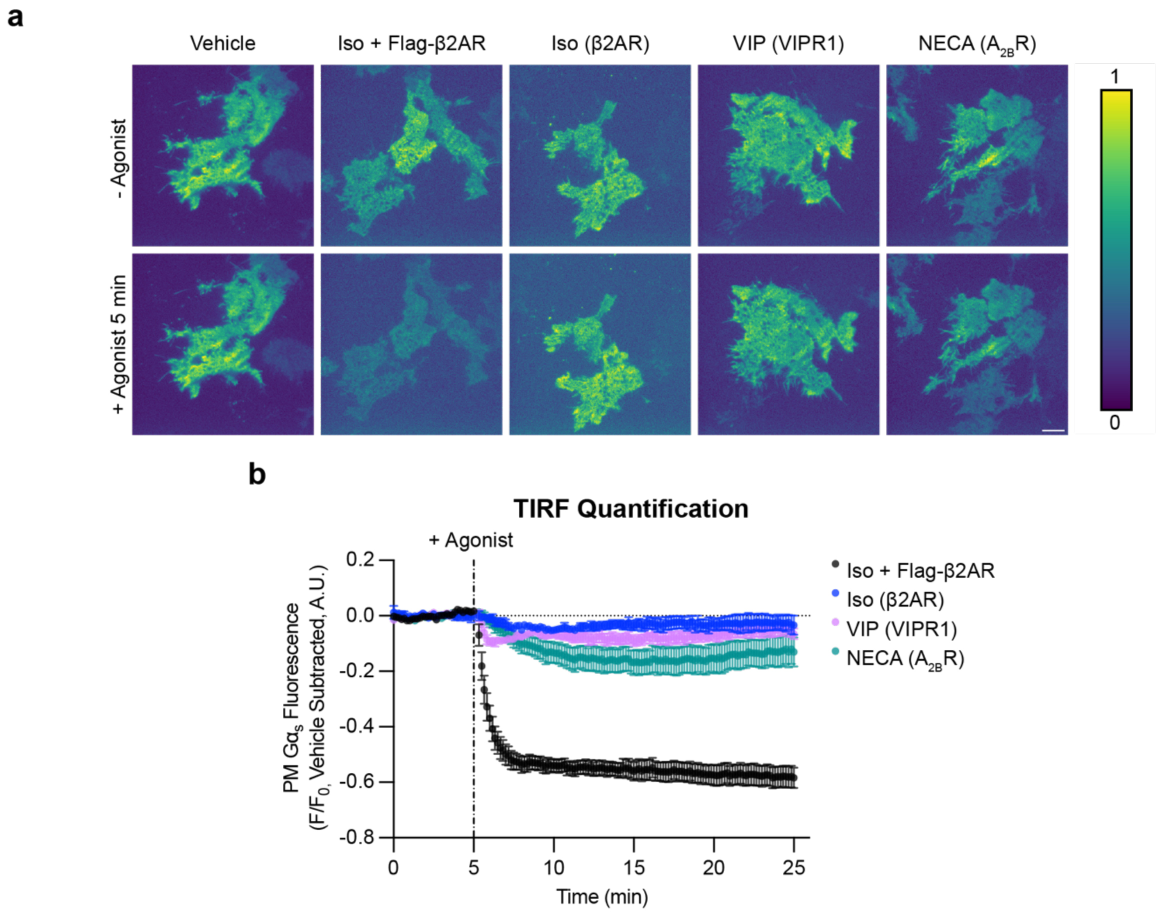


**Supplementary Figure 7: Endogenous GPCR activation triggers Gα$_s$ redistribution in an independent clone of EGFP-Gα$_s$ KO rescue cells. a)** Representative stills from time-lapse TIRF microscopy of Gα$_s$ KO2 + EGFP-Gα$_s$ rescue cells stably expressing EGFP-Gα$_s$ before or after 5 minutes of agonist treatment to activate endogenously expressed GPCRs (β2AR, VIPR1, $A_{2B}R$). Images are represented as heat maps normalized to t = 3 minutes before drug addition for each individual movie. Cells were treated with either vehicle, Iso (1 μM, ± overexpression of Flag-β2AR), VIP (1 μM), or NECA (20 μM). Scale bar = 10 μm. **b)** Quantification of Gα$_s$ fluorescence from TIRF movies depicted in a). The average of vehicle control movies (n = 4) at each time point was subtracted before plotting data. n = 4 movies from 4 experiments. Data are represented as mean ± S.E.M. of individual movies. Significance determined by repeated measures 2-way ANOVA with Dunnett's multiple comparisons test (see Supplementary Table 9 for p values).

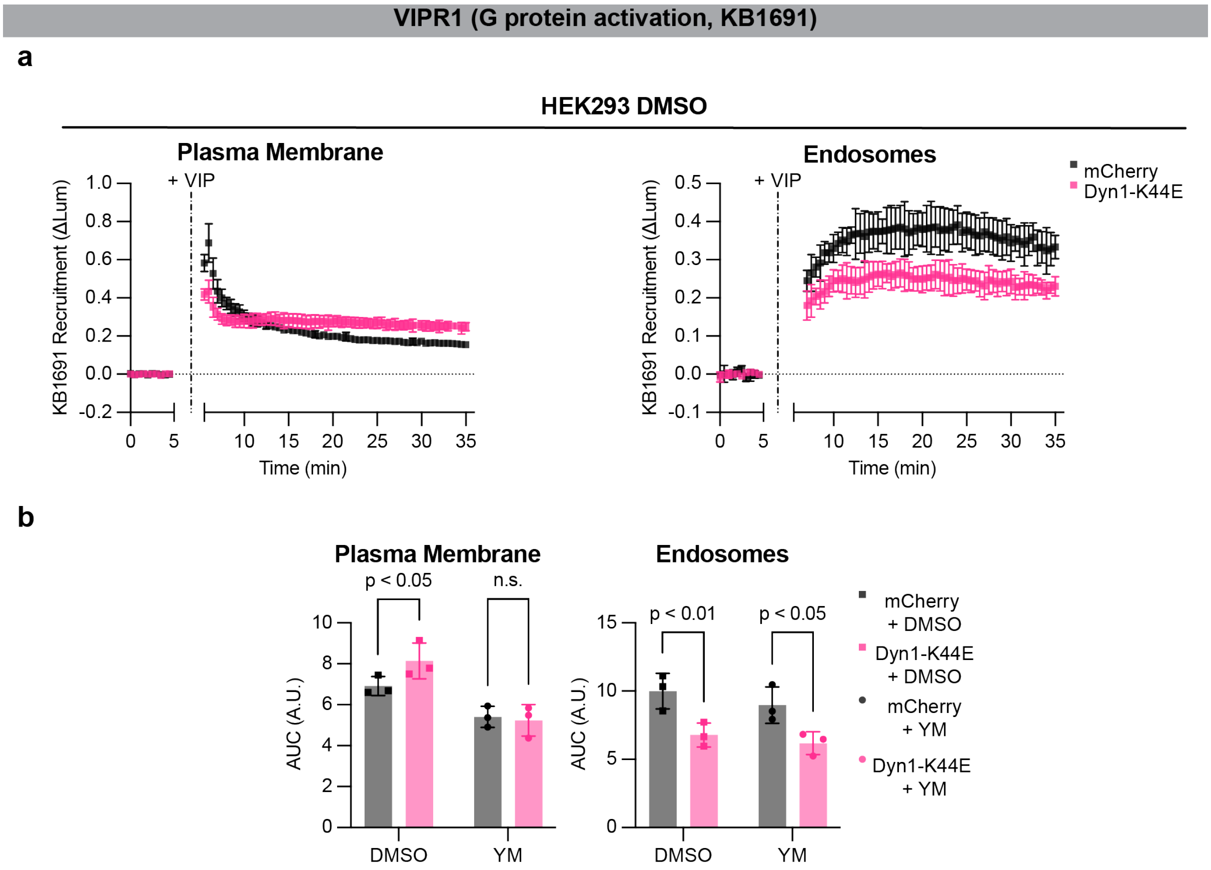


**Supplementary Figure 8: VIPR1 internalization is required for G protein activation at endosomes. a)** Left: DMSO control curves for KB1691 plasma membrane NanoBit bystander assay data shown in Figure 5e. Right: DMSO control curves for KB1691 plasma membrane NanoBit bystander assay data shown in Figure 5f. HEK293 cells expressing Halo-VIPR1 were pretreated with DMSO (0.1 %, 30 minutes) and VIP (1 μM) was added after 5 minutes. Cells were prepared in parallel with cells pretreated with YM-254890 (Fig. 5ef) and data were collected in the same plate. **b)** Area under the curve of NanoBit data in panel a (DMSO) and Figure 5ef (YM). YM bars are reproduced from Figure 5ef. Significance determined by repeated measures 2-way ANOVA with Sidak's multiple comparisons test (see Supplementary Table 12). Data are shown as mean ± S.D. of 3 independent experiments.

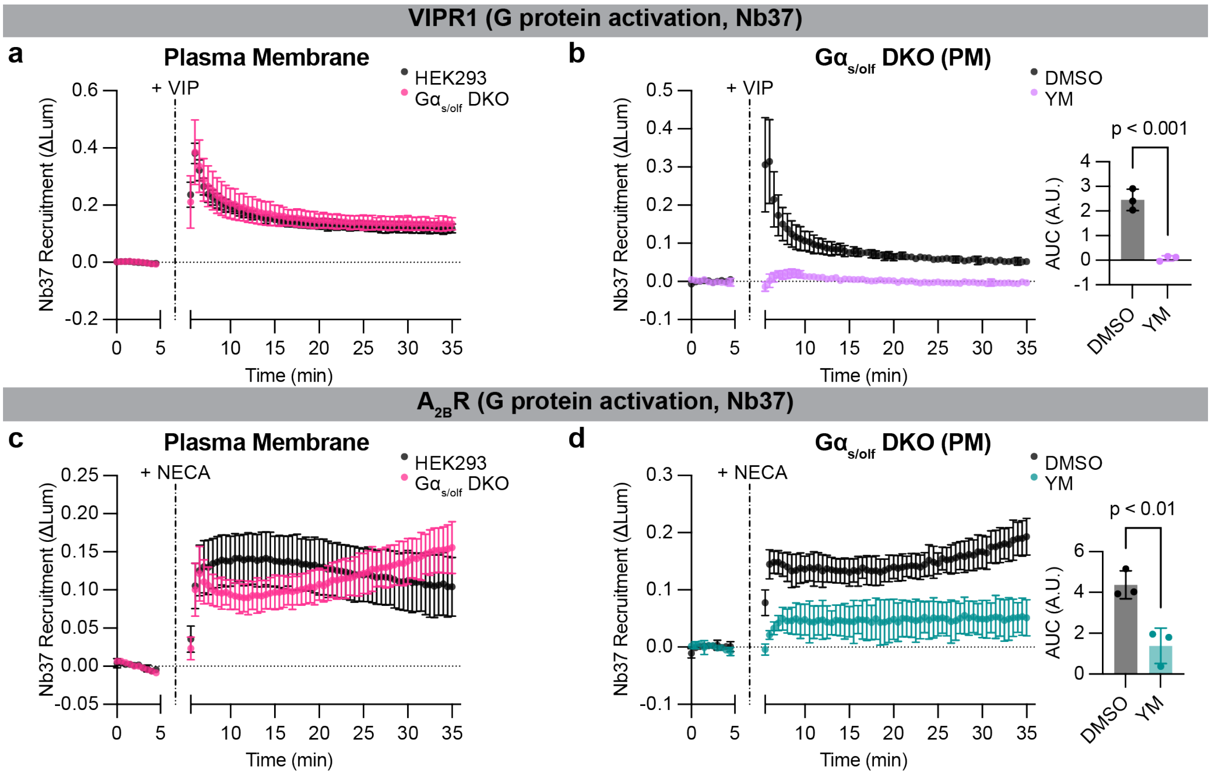


**Supplementary Figure 9: Nb37 detects G$\alpha_s$-independent, YM-sensitive signal. a)** NanoBit bystander assay showing recruitment of Nb37 to the plasma membrane in both HEK293 parental cells and G$\alpha_{s/olf}$ DKO cells expressing Halo-VIPR1. VIP (1 μM) was added after 5 minutes. **b)** Left: NanoBit bystander assay showing recruitment of Nb37 to the plasma membrane in G$\alpha_{s/olf}$ DKO cells expressing Halo-VIPR1 and pretreated with either DMSO (0.1 %) or YM-254890 (1 μM, 30 minutes). Right: AUC of time course. VIP (1 μM) was added after 5 minutes. Significance determined by unpaired, two-tailed t test. **c)** NanoBit bystander assay showing recruitment of Nb37 to the plasma membrane in both HEK293 parental cells and G$\alpha_{s/olf}$ DKO cells expressing Halo-$A_{2B}$R. NECA (100 μM) was added after 5 minutes. **d)** Left: NanoBit bystander assay showing recruitment of Nb37 to the plasma membrane in G$\alpha_{s/olf}$ DKO cells expressing Halo-$A_{2B}$R and pretreated with either DMSO (0.1 %) or YM-254890 (1 μM, 30 minutes). Right: AUC of time course. NECA (100 μM) was added after 5 minutes. Significance determined by unpaired, two-tailed t test. Data are shown as mean ± S.D. of least 3 independent experiments.
